# A unique and evolutionarily conserved retinal interneuron relays rod and cone input to the inner plexiform layer

**DOI:** 10.1101/2020.05.16.100008

**Authors:** Brent K. Young, Charu Ramakrishnan, Tushar Ganjawala, Yumei Li, Sangbae Kim, Ping Wang, Rui Chen, Karl Deisseroth, Ning Tian

## Abstract

Neurons in the CNS are distinguished from each other by their morphology, the types of the neurotransmitter they release, their synaptic connections, and their genetic profiles. While attempting to characterize the retinal bipolar cell (BC) input to retinal ganglion cells (RGCs), we discovered a previously undescribed type of interneuron in mice and primates. This interneuron shares some morphological, physiological, and molecular features with traditional BCs, such as having dendrites that ramify in the outer plexiform layer (OPL) and axons that ramify in the inner plexiform layer (IPL) to relay visual signals from photoreceptors to inner retinal neurons. It also shares some features with amacrine cells, particularly Aii amacrine cells, such as their axonal morphology and possibly the release of the inhibitory neurotransmitter glycine, along with the expression of some amacrine cell specific markers. Thus, we unveil an unrecognized type of interneuron, which may play unique roles in vision.

**Significance Statement:** Cell types are the building blocks upon which neural circuitry is based. In the retina, it is widely believed that all neuronal types have been identified. We describe a cell type, which we call the Campana cell, that does not fit into the conventional neuronal retina categories but is evolutionarily conserved. Unlike retinal bipolar cells, the Campana cell receives synaptic input from both rods and cones, has broad axonal ramifications, and may release an inhibitory neurotransmitter. Unlike retinal amacrine cells, the Campana cell receives direct photoreceptor input has bipolar-like ribbon synapses. With this discovery, we open the possibility for new forms of visual processing in the retina.

## Introduction

Photons entering the eye are detected by photoreceptors and processed through a set of function-specific signal pathways in the retina. The morphological basis of these pathways are the synaptic connections among five major classes of retinal neurons: photoreceptors, horizontal cells (HCs), bipolar cells (BCs), amacrine cells (ACs) and retinal ganglion cells (RGCs) (Gollisch and Meister, 2010; Asari and Meister, 2012; Masland, 2012). Two fundamental features of the visual signal processing in the retina are the functional separation of scotopic and photopic vision (Bloomfield and Völgyi, 2009; Dunn and Wong, 2014; Lamb, 2016) and the segregation of increment and decrement luminance signals into ON and OFF pathways (Wässle et al., 2009; Masland, 2012).

The functional separation of scotopic and photopic visions starts at rods and cones and remains separated at BCs due to specific synaptic connections from rods to rod BCs and cones to cone BCs (Azeredo da Silveira and Roska, 2011; Masland, 2012; Helmstaedter et al., 2013; Dunn and Wong, 2014). The segregation of increment and decrement luminance signals starts at BCs where glutamate released from cones activates ionotropic glutamate receptors on the OFF cone BCs resulting in depolarization (DeVries, 2000), while glutamate activates a metabotropic glutamate receptor, mGluR6, on the rod BCs and ON cone BCs resulting in hyperpolarization of these BCs (Nawy and Jahr, 1990, 1991; Shiells and Falk, 1990). This sign reversing and non-reversing action of glutamate on the ON and OFF BCs separates the increment and decrement luminance signals into ON and OFF pathways which remain segregated throughout the visual system (Pandarinath et al., 2010; Azeredo da Silveira and Roska, 2011; Dhande et al., 2015; Eiber et al., 2018). BCs are the only interneuron to relay visual signals from photoreceptors to RGCs, and they are the primary excitatory driver for RGCs (Saszik and DeVries, 2012; Lindstrom et al., 2014; Behrens et al., 2016; Shekhar et al., 2016). All BCs have a dendrite in the OPL and axons in the IPL, except for a recently identified monopolar BC (Santina et al., 2016). Additionally, it is commonly assumed that all BC types have been identified (Wässle et al., 2009; Baden et al., 2013a; Shekhar et al., 2016).

Another type of interneuron, ACs, do not receive direct synaptic input from photoreceptors, but instead from BCs and other ACs (Dowling and Boycott, 1966; Masland, 2012; Kim et al., 2014; Graydon et al., 2018). Most ACs act as inhibitory interneurons, through the release of GABA and glycine, except for a few glutamatergic and cholinergic types (O’Malley et al., 1992; MacNeil and Masland, 1998; Masland, 2001, 2012; Johnson et al., 2004; Keeley and Reese, 2010; Lee et al., 2010; Wei et al., 2011; Euler et al., 2014; Santina et al., 2016). Additionally, a small fraction of interplexiform ACs release dopamine as their neurotransmitters and conducting synaptic signals from the inner retina to the outer retina (Knapp and Dowling, 1987; Witkovsky, 2004; Jackson et al., 2012; Sankaran et al., 2018).

In this study, we identified a previously undescribed type of retinal interneuron. This interneuron shares some fundamental morphological, physiological, and molecular features with traditional BCs, such as having neurites that ramify in the OPL to synapse with photoreceptors and having neurites that ramify in the IPL to synapse with inner retinal neurons. Additionally, these cells express several common AC markers, and their neurites ramify throughout the entire IPL like Aii-ACs to signal neurons in both ON and OFF IPL. These interneurons may also release the inhibitory neurotransmitter glycine in both the inner and outer retina in addition to releasing glutamate in the IPL. These findings reveal an atypical retinal interneuron, which may play a unique role in visual processing.

## Materials and methods

### Animals

Wild type C57BL/6 mice and CreER-JamB:FRT-EGFP double transgenic mice with a C57BL/6 background of either sex aged 2-3 months were used in this study. CreER-JamB mice were generously given to us by Dr. Joshua Sanes, Harvard University (Buffelli et al., 2003; Kim et al., 2010). CreER-JamB:FRT-EGFP mice were generated by breeding CreER:JamB mice (obtained from Dr. Joshua Sanes, Harvard University) (Kim et al., 2008, 2010) with FRT-EGFP mice (MMRRC-032038-JAX) which harbor the R26R CAG-boosted EGFP (RCE) reporter allele with a FRT-flanked STOP cassette upstream of the enhanced green fluorescent protein (EGFP) gene (Sousa et al., 2009).

Primate retinal tissue was collected after the animals had been euthanized and transcardially perfused with saline followed by 4% PFA for 15-20 minutes.

All animal procedures and care were performed following protocols approved by the IACUC of the University of Utah in compliance with PHS guidelines and with those prescribed by the Association for Research in Vision and Ophthalmology (ARVO).

### Virus Injections

Four AAV viral vectors were used in this study. The AAV-EF1a-DIO mCherry-IRES-WGA-Flpo viral expression cassette was cloned in Dr. Karl Deisseroth’s lab (Stanford University). It was tested in rat hippocampal neurons for specificity and the absence of genetic leakage. It was used to express WGA-Flpo fused protein and mCherry in RGCs in a Cre-dependent manner in this study. This oversized genome was packaged into AAV serotype 2 capsid by UNC.

The AAV2-CAG-ChR2-GFP-Na1.6 viral expression cassette was previously described (Wu et al., 2011) and was used to express GFP in Aii-ACs and Campana cells in this study. Briefly, the construct was made by the addition of a 27 aa ankyrin binding domain from Na-v1.6 (NavII-III: 5’-TVRVPIAVGESDFENLNTEDVSSESDP-3’) (Garrido et al., 2003) to the 3’ end of ChR2-GFP fusion construct (Bi et al., 2006). The AAV2-CAG-GCaMP6m viral expression cassette was made by swapping out the ChR2-GFP with GCaMP6m. The DNA of GCaMP6m was synthesized (GenScript, Piscataway, NJ) according to the reported amino acid sequence (Chen et al., 2013). For both AAV2-CAG-ChR2-GFP-Na1.6 and AAV2-CAG-GCaMP6m viral vectors, the ChR2-GFP and GCaMP6m plasmids were packaged into AAV serotype 2 capsid by Virovek (Hayward, CA, USA). Additionally, the AAV2.7m8-Y444F-mGluR500p-mCherry virus was provided by Dr. Zhuo-Hau Pan’s lab (Wayne State University) and was used to label rod bipolar cells (Lu et al., 2016).

Viruses were injected intraocularly into the mice to induce in vivo gene expression at ages P11-P30 using the procedure described previously in detail (Xu et al., 2010). For intraocular injection, the mice were anesthetized with Isoflurane (2-4% Isoflurane mixed with room air delivered in a rate between 0.8 and 0.9 L/min, Kent Scientific, Torrington, CT) through a mouse gas anesthesia head holder (David KOPF Instruments, Tujunga, CA, United States). 0.5% proparacaine hydrochloride ophthalmic solution was locally applied to each eye. Glass needles made from borosilicate glass using a Brown-Flaming horizontal puller (P-1000 puller, Sutter Instruments) with a fine tip (about 10-15 μm diameter) were used for injection. The glass needles were mounted on a Nano-injection system (Nanoject II, Drummond Scientific Company, Broomall, PA, United States), which could precisely control the amount of injected solution at the nanoliter level, and the Nanoject was mounted on a micromanipulator (M3301R, World Precision Instruments). For each eye, 1-2µl of the solution was injected over 3-5 minutes. After the injection, the eyes were covered with ophthalmic ointment, and the mice were placed in a clean cage sitting on a heated water blanket. The temperature of the water blanket was set at 33°C. Mice were continuously monitored until they completely recovered, and then they were returned to their home cages. The procedures for anesthesia and intraocular injection fit the procedures approved by the IACUC of the University of Utah.

### WGA-Flpo technique

We use IP injection of Tamoxifen to induce CreER activation. The efficiency and duration of Tamoxifen-induced CreER activation depends significantly upon the strain of mice, type of promotor for the transgene, the dosage of Tamoxifen administered, and the age of the mice receiving the Tamoxifen (Reinert et al., 2012). Accordingly, we have optimized the injection of Tamoxifen and viral vector paradigm for age, strains, and dosage. In order to activate CreER in a large population of RGCs, we injected Tamoxifen at a young age (P7-P14), but in order to prevent non-specific transduction and expression, we found that the virus needed to be injected slightly later. Early studies showed that Cre remains active for a day (Brocard et al., 1997), but this was in adult mice and used approximately 15% of the Tamoxifen concentration that we use (150mg/kg). More recent studies have shown that Tamoxifen remains in the system and available for CreER activation for at least 4 weeks after injection and is 46% efficient a week after injection using a concentration similar to ours (Reinert et al., 2012). Therefore, we developed a protocol that maximizes the specificity and efficiency of gene expression. In addition to all of this, we performed a time course of cellular activation of CreER in J-RGCs and found that they can be activated by Tamoxifen up to P40. In accordance with this protocol, we treated mice with Tamoxifen (150mg/kg) one week before the injection of the AAV2-EF1a-DIO mCherry-IRES-WGA-Flpo viral vector into CreER-JamB:FRT-EGFP mice.

The WGA-Flpo approach utilizes a dual DNA recombinase strategy to express WGA (wheat germ agglutinin)-Flpo recombinase fused protein in Cre-positive RGCs using a Cre-dependent mCherry:IRES-WGA-Flpo viral vector (WGA-Flpo) on Cre:FRT-EGFP double transgenic mice. WGA is a plant lectin and a commonly used transcellular tracer. Once WGA is introduced into neurons, it can be transported transcellularly by exocytosis and is taken up by the pre/post-synaptic neurons through endocytosis (Shibata et al., 1986; Ilinsky and Kultas-ilinsky, 1990; Tabuchi et al., 2000; Yoshihara, 2002; Damak et al., 2008; Gradinaru et al., 2010; Libbrecht et al., 2017). WGA binds to N-acetylglucosamine and sialic acid carbohydrate residues on the cell surface and is then transported through cell membranes in cytoskeleton-dependent clathrin and caveolae-mediated mechanisms (Broadwell and Balin, 1985; Tabuchi et al., 2000; Yoshihara, 2002; Gao et al., 2008; Libbrecht et al., 2017). WGA has been used for transcellular tracing in many areas throughout the CNS, including the retina, and WGA fused to Cre, was also found to be transported across synapses (Tabuchi et al., 2000; Braz et al., 2002; Gradinaru et al., 2010; Reeber et al., 2011; Ieki et al., 2013; Libbrecht et al., 2017).

We injected the WGA-Flpo vector into the eyes of CreER-JamB:FRT-EGFP mice. These mice express Cre DNA recombinase in a function-specific RGC type, the J-RGCs (Kim et al., 2008), and harbor the R26R CAG-boosted EGFP reporter allele with an *FRT*-flanked STOP cassette in all somatic cells (Sousa et al., 2009). J-RGCs are OFF-RGCs that have been found to activate in response to a variety of properties (Kim et al., 2008; Joesch and Meister, 2016; Nath and Schwartz, 2017). The WGA-Flpo vector transduces Cre-positive J-RGCs in these mice and expresses both mCherry and WGA-Flpo fused proteins. When WGA-Flpo fused protein is transported into neurons synaptically connected to the transduced Cre-positive J-RGCs, mEGFP is expressed in these cells, which used to illustrate the cell morphology.

### Immunohistochemistry

The procedures for fluorescent immunolabeling of retinal neurons on retinal whole-mounts and cross-section preparations have been described previously in detail (Xu et al., 2010). In brief, mice were euthanized with 100% CO2, followed by cervical dislocation. Eyes were enucleated, the cornea was slit, and then the eyeballs were fixed in 4% paraformaldehyde (PFA) in 0.01M phosphate-buffered saline (PBS; pH 7.4) for 1-1.5 hours at room temperature (RT). Following fixation, the retinas were isolated and washed 20 min × 4 in 0.01M PBS at RT. Retinas were then blocked (5% normal donkey serum, 0.5% Triton X-100 in 0.01M PBS) for 1.5 hours at RT. Next, retinas were incubated in primary antibody diluted in blocking solution for 6-7 days, washed 20 min × 4 in 0.01M PBS at RT, and then incubated in secondary antibody for 1-2 days. Finally, retinas were flat-mounted with spacer (Grace Bio-Labs) in H-1000 Vectashield (Vector Laboratories).

For cryosections, the cornea of enucleated eyes was slit and fixed 1.5-2 hours in 4% PFA at RT. In one experiment to label the mGluR6 in the retina, the eyes were fixed for only 15 minutes (Morgans et al., 2006). Eyes were then cryoprotected in 10% sucrose (2 hours at RT), 20% sucrose (overnight at 4°C), and then 30% sucrose (overnight at 4°C). Cryoprotection of primate retinal tissue was similar, except for the eye globes were kept in 10% sucrose for 24 hours, 20% sucrose for 2 days, and 30% sucrose for 1-2 weeks. Following cryoprotection, eyes were mounted in OCT Compound (Fisher Scientific) and frozen on dry ice. Sections were cut at a thickness of 30-50 µm. Sections were dried at RT for 10 mins, hydrated in 0.01M PBS for 10 mins at RT, then blocked in blocking solution for 1-2 hours at RT. The primary antibody was diluted in blocking solution and placed on sections overnight at 4°C or RT depending on the primary antibodies used. Sections were washed 3×15mins in 0.01M PBS at RT, and then the secondary antibody was placed on the sections for 2 hours at RT. Finally, sections were washed 3×15mins in 0.01M PBS at RT, stained with DAPI (1µg/ml) for 30 mins, washed for 30 seconds, and then mounted in Vectashield. All primary antibodies were validated through the use of a control staining with only the secondary antibody used. The details of all primary and secondary antibodies used in this study, the source, and concentration are listed in Table 1.

**Table 1.** Antibodies

| Antibody | Host | Concentration | Supplier | Catalog # | Reference |
| --- | --- | --- | --- | --- | --- |
| CaB5/CaBP5 | Rabbit | 1:500 | Haeseleer |  | (Haeseleer et al., 2000) |
| Calretinin | Goat | 1:1000 | EMD Millipore | AB1550 |  |
| Chat | Goat | 1:100 | Chemicon | AB1440 |  |
| Cone Arrestin | Rabbit | 1:1000 | EMD Millipore | AB15282 |  |
| CtBP2 | Mouse | 1:1000 | BD Biosciences | 612044 |  |
| Glutamic Acid<br>Decarboxylase | Mouse | 1:50 | Developmental<br>Studies Hybridoma<br>Bank | GAD-6 |  |
| GAD67 | Rabbit | 1:500 | Chemicon | AB108 |  |
| GFP-FITC | Goat | 1:500 | GeneTex | GTX26662 |  |
| GFP-488 | Rabbit | 1:500 | ThermoFisher | A21311 |  |
| GlyT1 | Goat | 1:10000 | EMD Millipore | AB1770 | Botir T.<br>Sagdullaev |
| GlyRa1+2 | Rabbit | 1:750 | Abcam | AB23809 |  |
| mGluR6 | Sheep | 1:100 | Morgan Lab |  | (Morgans<br>et al.,<br>2006) |
| Otx2 | Goat | 1:1000 | R&D Systems | AF1979-SP |  |
| Pax6 | Rabbit | 1:500 | Biolegend | 901301 |  |
| PKC | Mouse | 1:100 | Abcam | ab31 |  |
| REEP6 | Rabbit | 1:1000 | Swaroop Lab |  | (Hao et al.,<br>2014) |
| RFP | Rabbit | 1:1000 | Rockland Antibodies | 600-401-379 |  |
| SV2 | Mouse | 1:50 | Developmental<br>Studies Hybridoma<br>Bank | SV2 |  |
| VGAT | Rabbit | 1:500 | EMD Millipore | AB5062P |  |
| VGlut1 | Guinea<br>Pig | 1:500 | EMD Millipore | AB5905 |  |
| VSX2/Chx10 | Sheep | 1:300 | Exalpha biologicals<br>INC | X1180P |  |
| WGA | Goat | 1:1000 | Vector Laboratories | AS-2024 |  |
| Goat:Alexa488 | Donkey | 1:400 | Jackson<br>ImmunoResearch | 705-545-147 |  |
| Goat:Cy3 | Donkey | 1:400 | Jackson<br>ImmunoResearch | 705-165-147 |  |
| Goat:Cy5 | Donkey | 1:400 | Jackson<br>ImmunoResearch | 705-175-147 |  |
| Guinea pig:Cy3 | Donkey | 1:400 | Jackson<br>ImmunoResearch | 706-165-148 |  |
| Guinea pig:Cy5 | Donkey | 1:400 | Jackson<br>ImmunoResearch | 706-175-148 |  |
| Mouse:Cy3 | Donkey | 1:400 | Jackson<br>ImmunoResearch | 715-165-151 |  |
| Mouse:Cy5 | Donkey | 1:400 | Jackson<br>ImmunoResearch | 715-175-150 |  |
| Sheep:Cy3 | Donkey | 1:400 | Jackson<br>ImmunoResearch | 713-165-147 |  |
| Rabbit:Alexa405 | Donkey | 1:400 | Abcam | ab175651 |  |
| Rabbit:Cy3 | Donkey | 1:400 | Jackson<br>ImmunoResearch | 711-165-152 |  |
| Rabbit:Cy5 | Donkey | 1:400 | Jackson<br>ImmunoResearch | 711-175-152 |  |

### Single Cell sequencing

Single cells were collected for RNA profile from wild type C57BL/6 mice intraocularly injected either with AAV2-CAG-ChR2-GFP-Na1.6 (for Aii-ACs and Campana cells), or AAV2.7m8-Y444F-mGluR500p-mCherry (for rod BCs). Around 3-5 weeks after viral vector injection, mice were euthanized with 100% CO2; eyes were enucleated, retinas were isolated in RNase free HEPES buffer (137 mM NaCl, 5 mM KCl, 2.5 mM CaCl2 (2H20), 1 mM MgCl (6H20), 10.5 mM HEPES-NaOH, 22 mM Glucose, pH adjusted to 7.4 with NaOH), mounted on filter paper, and cut into 200 µm sections. Retina sections were placed in a custom build chamber and perfused with oxygenated RNase free HEPES buffer. Aii-ACs, Campana cells, and rod BCs were identified based on GFP expression and morphology. A glass needle (Garner Glass Company) was front-loaded with ∼1 µl RNase free 0.01M PBS (Thermo Fisher Scientific), contacted to the identified Aii-ACs, Campana cells, or rod BCs, and the contents of the soma of the cell were suctioned into the needle (Figure S8). We collected the cellular contents of 5-10 Campana cells, Aii-ACs, and rod BCs fluorescently labeled by AAV2-GFP or AAV2.7m8-mGluR6-mCherry (AAV2.7-mCherry) (Lu et al., 2016) viral vectors (Figure 8). The contents were then placed in a 3 µl droplet of RNase free 0.01M PBS, which was flash-frozen in liquid nitrogen and stored at −80° C. RNAseq library was generated according to SMART-seq v4 Ultra-low input RNA kit (Takara Clontech). In brief, isolated cells were lysed in lysis buffer, 3-SMART-seq CDS primer II and V4 oligonucleotide were added for first stranded cDNA synthesis. cDNA was amplified using PCR Primer II A, and subsequently purified using Ampure XP beads (Beckman). Illumina library was prepared using the Nextera XT DNA library preparation kit (Illumina) and sequenced using Illumina Novaseq (Daines et al., 2011; Zhao et al., 2016). The sequencing analysis was done blind to the cell type. We sequenced 34,118,309 reads from these cells and 20,289 actively expressed transcripts were detected. From these actively expressed transcripts, 6913 are uniquely expressed by the 3 types of neurons (6301 by Aii-ACs, 415 by Campana cells, and 197 by rod BCs). Next, we used FPKM (Fragments Per Kilobase of transcript per Million mapped reads) profile for a comparison between the samples after removing unexpressed transcripts across all of the samples (13,237 transcripts) and performing quantile normalization (limma package, R).

### Image acquisition and analysis

Whole-mount retinas were imaged using a two-photon microscope (Bruker), with a 40× 1.0 NA Zeiss water immersion objective. Retina sections were imaged using an LSM 700 or LSM 800 confocal microscope (Zeiss) with a 40x 1.2 NA water immersion, or 63×1.4 NA oil immersion Zeiss objectives. Pre/post-synaptic marker imaging was done with the 63× Zeiss objective with a maximum theoretical lateral resolution of 150-190 nm and an axial resolution of 550-720 nm (Smith, 2008). Verification of pre/post-synaptic marker localization was done with the 63× Zeiss objective at super-resolution using Zeiss airyscan that has a lateral resolution of 120 nm and axial resolution of 350 nm (Müller and Enderlein, 2010; York et al., 2012, 2013; Roth and Heintzmann, 2016).

BCs were identified based on their morphology, with the axonal ramification depth in the IPL providing separation between ON or OFF cone BCs, and then we used axonal morphology to differentiate the types (Tsukamoto and Omi, 2017; Wässle et al., 2009).

Maximum projections of imaging stacks, adjustments of brightness/contrast for individual channels, and single-plane images were generated using the Fiji plugin for ImageJ2 (Schindelin et al., 2012; Rueden et al., 2017). Image processing, video creation, 3D rendering, and data analysis were done using Imaris software (Bitplane), and video files were edited using Adobe Premiere Pro (Adobe). Measurements of axonal and dendritic field areas were made from flat-mount retinas. Convex polygon area measurements were made using the Fiji plugin for ImageJ2 (Schindelin et al., 2012; Rueden et al., 2017), and assumed a 2D field from the outer most points of either the dendrite (OPL) or axon (IPL). Nearest neighbor measurements were made using Fiji and each measurement was taken from the center of the soma. Random Nearest Neighbor calculation was made through the probability density *p*(*r*) formula:

*p(r)* = *k*2πλ*r* exp(-λπ*r*^2^)

where *k* is a normalization factor, λ is the cell density, and *r* is a given distance from an arbitrary point (Wässle and Riemann, 1978). The Gaussian fit for the Nearest Neighbor Distribution was made using Igor (Wavemetrics).

For the calculation of colocalization, a surface mask was made of both channels, and then colocalized objects were restricted to those objects that were fully enveloped within the cell (no more than 2 pixels, ≤100 µm, outside of the cell). As a negative control, we measured the colocalization of ribbon synapses inside of Aii-ACs using the same technique. This resulted in an average of 4.24 ± 0.56 ribbons per cell which is 6.5 times fewer than what we observed in Campana cells (unpaired t-test, p = 0.0036). It is unclear if the CtBP2 colocalization with Aii-ACs is real, or an imaging artifact. CtBP2 staining of ACs has not been previously reported, but can be observed in previous publications (Lee et al., 2015; Strettoi et al., 2018).

To measure the total number of close associations with cones and rods of each Campana cell, we verified each cell as a Campana cell by tracing and masking the GFP signal (Figure 4C). We only used cells that had their entire dendritic structure included in a single retinal section (40 µm in thickness). Figure 4D shows a 90-degree rotation of the entire dendritic plexus from the dash-lined box in Figure 4C. The synaptic connections between Campana cells and rods/cones were identified based on the close association of the dendritic terminals of Campana cells and the ribbons located inside the axonal terminals of rods or cones similar to previous work (Strettoi et al., 2018). To calculate rod and cone input, we measured the distance from the dendritic terminals of Campana cells to ribbon synapses of either rods or cones. Those ribbons that were ≤300 nm away were counted as synaptic partners since a similar distance can be measured from ribbons to ON cone BCs in EM sections (Sato et al., 2008; Katiyar et al., 2015).

### Two-photon calcium imaging and light stimulation

For ex vivo imaging of calcium transients, all mice were treated with intraocular injection of AAV2-CAG-GCaMP6m viral vector 3-5 weeks before recording. One day before recording, mice were dark-adapted for >12 hours. Retinas were dissected in carboxygenated (95% O2, 5% CO2) mouse ringers (124 mM NaCl, 2.5 mM KCl, 2 mM CaCl2, 2 mM MgCl2, 1.25 mM NaH2PO4, 26 mM NaHCO3, and 22 mM glucose, pH 7.4), under infrared illumination. Retinas were then flattened on nitrocellulose filter paper (Millipore Corp) before being moved to a custom-made perfusion chamber. The chamber was then perfused with carboxygenated mouse ringers heated so that a temperature of 32°C was maintained at the retina. A two-photon microscope (Bruker) equipped with a mode-locked Ti:Sapphire laser (MaiTai, Newport Spectra-Physics) was used to image calcium transients. To record from BCs, we used a Zeiss 40x 1 NA objective (Zeiss).

Consistent with previous reports (Baden et al., 2013c, 2016; Palczewska et al., 2014; Hsiang et al., 2017), the two-photon laser could elicit a light response from some neurons in the retina, including Campana cells (Figure 3—supplement figure 1A and 1C). This light response could be bleached with a sustained dim green background light (Figure 3— supplement figure 1B, or locally reduced through 30 seconds of continuous scanning (laser bleach) before presenting a 10 ms light stimulus (Figure 3—supplement figure 1D. However, cone BCs did not respond to the laser scanning of the two-photon (Figure 5E and 5L). Therefore, we interpret that the two-photon laser stimulates rods at our excitation wavelength (955 nm), and we use the laser bleach protocol to isolate the cone light responses of Campana cells. Due to the opsin sensitivity to the two-photon laser, all cone responses were recorded after 30 seconds of continuous laser scanning (Baden et al., 2013c, 2016; Hsiang et al., 2017). Pharmacological reagents were used at the following concentrations, AP4 50 µM, AP5 35 µM, and CNQX 10 µM.

The light stimulation for Campana cells and type 5 cone BCs consisted of 30 seconds of continuous imaging followed by a 10 ms light stimulus, immediately after which we imaged for 15-30 seconds. A 180-second wait followed this with no stimulus or imaging before proceeding to the next light intensity, after which we repeated the same protocol with higher light intensity. Imaging was done at 28-33 frames per second.

Mice express a single rod photopigment, rhodopsin, and two cone photopigments, M-opsin, and S-opsin. Since rhodopsin and M-opsin have peak sensitivity at 509-512 nm, while the S-opsin peaks at 360 nm we used a dim green light that produced 0.549 x 10^3^ photons/(um^2^ x sec) for the rod response (Jacobs et al., 1991, 2004; Sun et al., 1997; Fu and Yau, 2007). Full-field light stimulation was provided by a custom-built LED light array. There were 4 LEDs, each of green (peak wavelength: 520 nm) and UV (peak wavelength: 400 nm). Light intensity was measured by a Model 371 optical power meter (UDT Instruments), and the light spectrum was measured by a C-700 Spectromaster Color Meter (Sekonic).

## Results

### Newly discovered interneurons have unique morphological features

To identify the neurons synaptically connected to RGCs, we first used a transgenic/viral approach to transcellularly label presynaptic neurons of RGCs (WGA-Flpo technique; see Methods). Using this approach, we examined the types of BCs presynaptic to RGCs (retinal ganglion cells). In addition to observing rod BCs and cone BCs, we unexpectedly observed a type of interneuron, which has not been described previously. Due to the synaptic transfer of WGA-Flpo, and the possibly of two gap-junction coupled cells appearing as one, we then attempted to find a method to label these new cells *without* relying on synaptic transfer. We found that a previously published viral construct, AAV2-CAG-ChR2-GFP-Na1.6 (AAV2-GFP; Figure 1A) (Wu et al., 2011), was able to label a variety of different retinal cell types following intraocular injection (Figure 1B). These included type 5 cone BCs (Figure 1C), Aii-ACs (Figure 1E), and the new interneurons that we found using the WGA-Flpo technique (Figures 1D). We characterized the morphology of type 5 cone BCs through isolating the GFP signal and confirming that its axons ramify at the start of sublamina b (the ON sublamina) of the IPL (Figure 1C).

**Figure 1.**
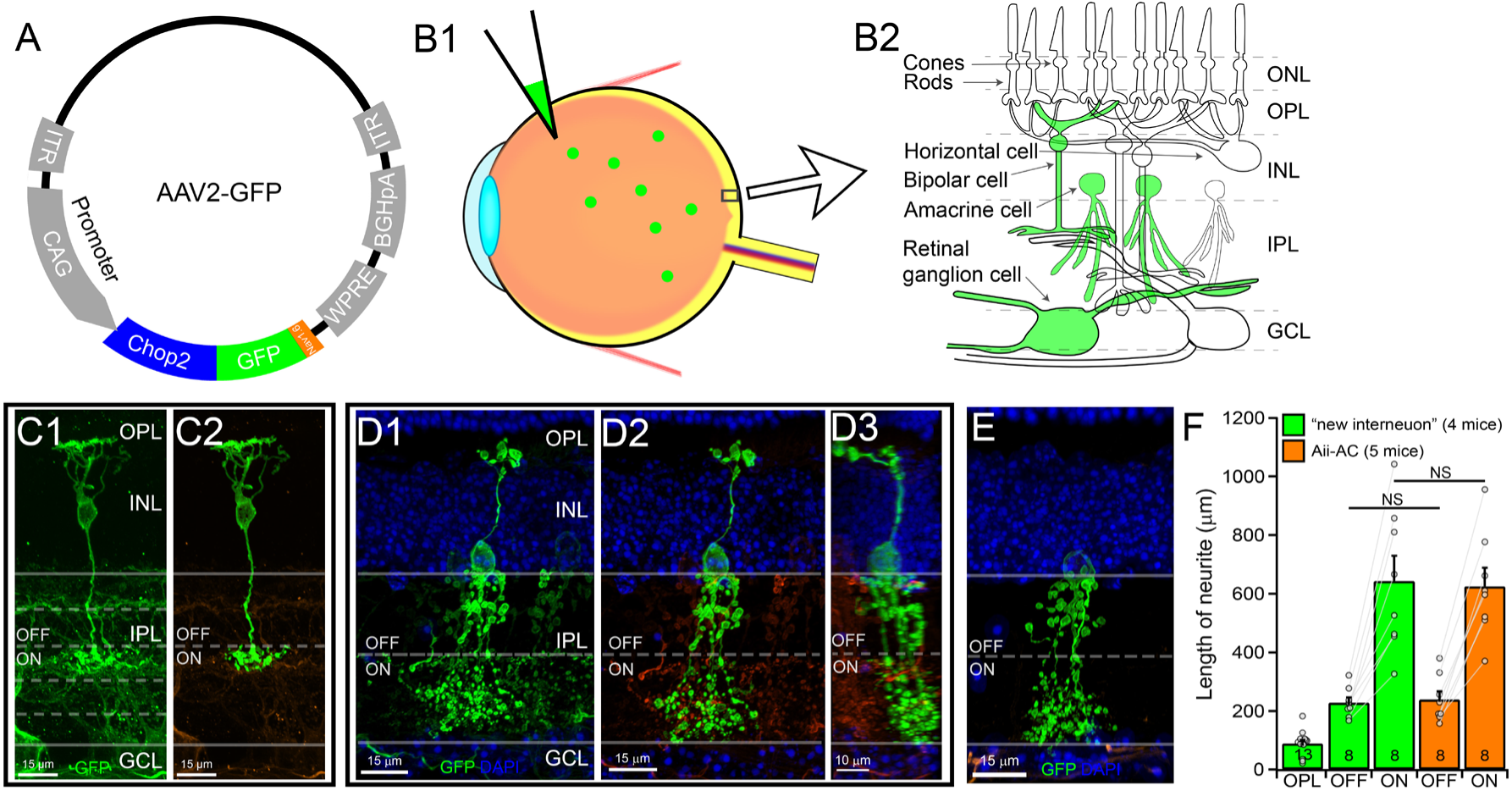
Newly discovered interneurons have unique morphological features. **A:** Structure of the AAV2-CAG-ChR2-GFP-Na1.6 viral cassette. **B:** Depiction of the use of the AAV2-CAG-ChR2-GFP-Na1.6 virus for intraocular injections (B1). Following intraocular injection, AAV2-CAG-ChR2-GFP-Na1.6 causes GFP expression (green) in a variety of cell types in the retina, but predominately infects Aii-ACs at a low viral titer (B2). **C:** Image of a type 5 cone BC labeled by AAV2-CAG-ChR2-GFP-Na1.6 (AAV2-GFP) vector in a thick (40 μm) retinal cross-section (C1, green), and the GFP within the cell masked from the background (orange) to reveal the morphology (C2). **D:** The newly identified interneuron labeled by AAV2-GFP (D1, green) and the same cell (green) masked and isolated from the background (orange, D2). D3 shows the same cell with a 90° rotation about the Y-axis. **E:** An Aii-AC labeled by AAV2-GFP shows similar axonal morphology to these newly identified cells. **F:** A comparison of the mean neurite length of the “new interneuron” (green) and Aii-ACs (orange) in the OFF (229 ± 18 µm “new interneuron” vs. 240 ± 28 µm Aii-AC, p = 0.7456) and ON (643 ± 86 µm “new interneuron” vs. 625 ± 63 µm Aii-AC, p = 0.8692) sublamina of IPL showed no significant difference between the mean length of the neurites (unpaired student t-test). The number in each column indicates the number of cells. Data are represented as mean + SEM. See also supplemental video 1 associated with this figure.

Similar to conventional BCs, these new interneurons have plexuses of neurites that ramify in both the OPL and IPL (Figures 1D). However, unlike conventional BCs that generally stratify their axonal terminals narrowly in either sublamina a (OFF) or sublamina b (ON) of the IPL (Wässle et al., 2009; Euler et al., 2014; Tsukamoto and Omi, 2017), this new interneuron has neurite ramifications throughout the IPL (Figures 1D). Additionally, there is a neurite that goes from the soma to the OPL (Figure 1D). This OPL projecting neurite was confirmed to be from this new interneuron, since there are *no other somas from other cells nearby*, even when the imaging is rotated (Figure 1D3 and Figure 1— supplemental video 1). It was striking to us how similar the IPL neurites of this new interneuron were to those of Aii-ACs. Therefore, we compared the length of the neurites of the “new interneuron” with those of Aii-AC, and we found that there was no significant difference between these new interneurons and Aii-ACs (OFF layer: 229 ± 18 µm “new interneuron” vs. 240 ± 28 µm Aii-AC, p = 0.7456, and ON layer: 643 ± 86 µm “new interneuron” vs. 625 ± 63 µm Aii-AC, p = 0.8692, student t-test; Figure 1F). However, Aii-ACs do not have a neurite that projects to the OPL like these “new interneurons” do (88.8 ± 11.9 µm; Figure 1F). Since this new interneuron looks similar to a hand bell, with a long neurite that travels to the OPL through in INL (the handle), and broad neuronal ramifications throughout the depth of the IPL (the bell), we call them “Campana cells.” *Campana* being Latin for “bell.”

### The Campana cell has some protein expression in common with classic BCs

Since the Campana cell has a neurite like a BC dendrite, we examined if they express synaptic proteins and synapse with photoreceptors through this neurite just as BCs do. In the vertebrate retina, both rod BCs and ON cone BCs express mGluR6 at their dendritic terminals (Nakajima et al., 1993; Nomura et al., 1994), while OFF cone BCs express both KA (kainic acid) and AMPA (α-amino-3-hydroxy-5-methyl-4-isoxazolepropionic) receptors (DeVries, 2000). Accordingly, we examined if Campana cells express mGluR6 (Morgans et al., 2006). Figure 2A (see also Figure 2—supplemental video 1) shows one of the Campana cells co-labeled by mGluR6, CtBP2, and cone arrestin (cone photoreceptors) (Liu et al., 2012). Super-resolution imaging of this same cell demonstrated close proximity between a ribbon synapse in a cone pedicle and mGluR6 in the neurite in the OPL (Figure 2B). A 3D colocalization analysis confirmed this proximity (Figures 2B4 and Figure 2— supplemental video 3). Because the neurite in the OPL of the Campana cells seems to be postsynaptic to photoreceptors with mGluR6 receptors, we hereafter refer to this as a dendrite.

**Figure 2.**
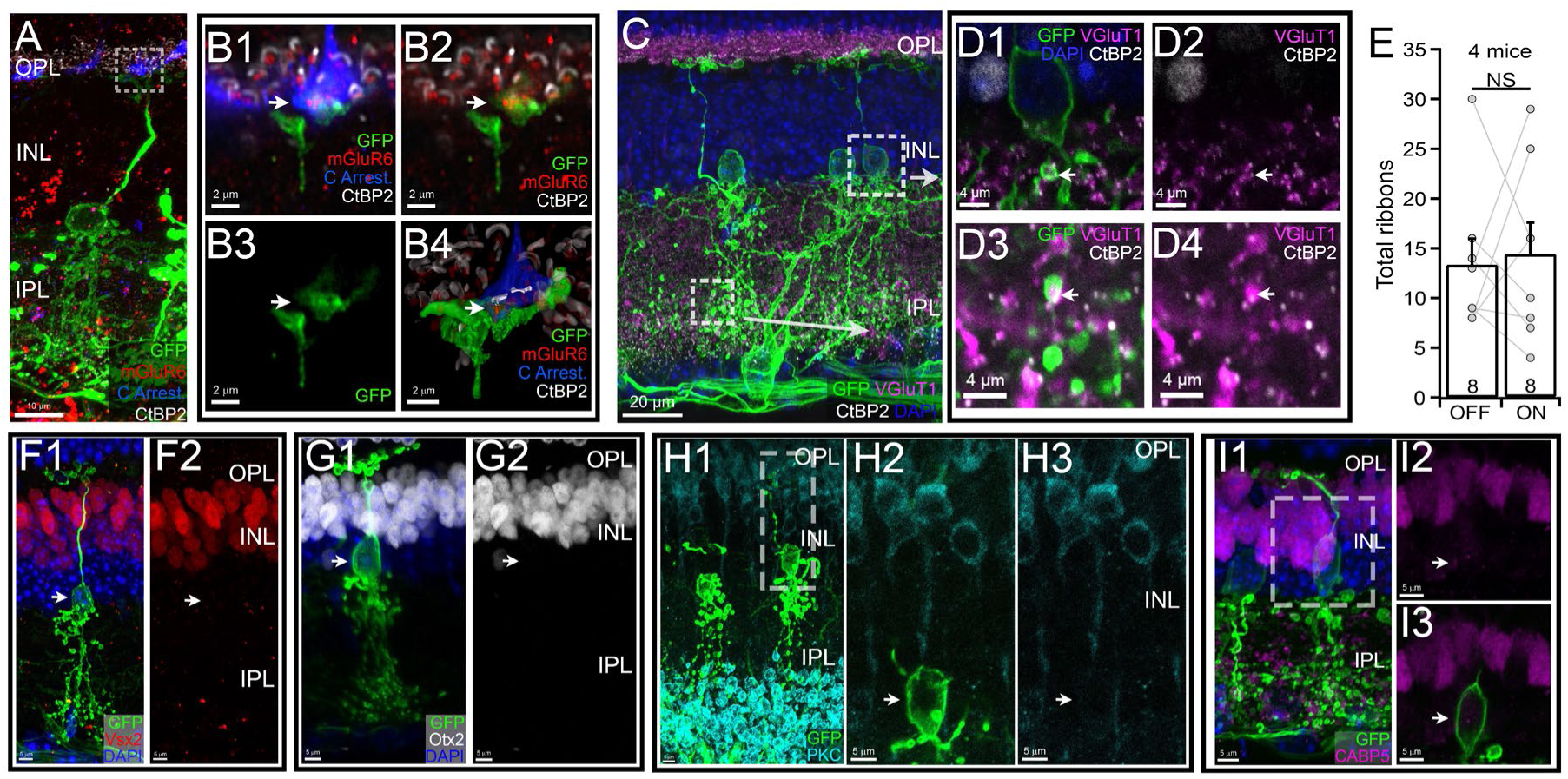
The Campana cell expresses some proteins in common with classic BCs. A: An AAV2-GFP labeled Campana cell (green) co-labeled by anti-cone arrestin (blue), anti-CtBP2 (white), and anti-mGluR6 (red) antibodies. **B:** A super-resolution single optic section of the area indicated by the box in 3A shows the dendritic terminal of the Campana cell (green, B1-B3) and anti-mGluR6 staining (red, B1 and B2). Anti-mGluR6 antibody stained puncta (red) are located inside the dendritic terminal of the Campana cell (green) and are closely associated with anti-CtBP2 labeling (white, B1 and B2) inside a cone terminal (blue, B1). Three-dimensional colocalization analysis confirms that mGluR6 is inside the Campana cell dendrite and is closely associated (≤300 nm away) with the ribbon synapse of a cone (B4). Arrows indicate the pre/post-synaptic structures. **C:** Two Campana cells side by side labeled with AAV2-GFP (green), and co-labeled with anti-CtBP2 (white), and anti-VGluT1 (magenta) antibodies. **D:** Magnified single optic section view of the two areas indicated by the boxes in 2C shows that ribbon synapses (anti-CtBP2, white), and anti-VGluT1 staining (magenta) are inside the axonal terminals (green, indicated by white arrow) in both ON (D3 and D4) and OFF (D1 and D2) layers of IPL. **E:** A comparison of the mean number of ribbon synapses of Campana cells localized within the OFF (13.38 ± 7.4) and ON (14.38 ± 8.9) sublamina shows the difference is not significant (p = 0.81, paired student t-test). Each dot represents the value of a cell. The number in each column indicates the number of cells. Data are represented as mean + SEM. **F:** A maximum projection image of an AAV2-GFP labeled Campana cell (green, F1) co-labeled with anti-Vsx2 antibody (red, F1, and F2) shows that the Campana cell is Vsx2-negative (F2). The arrow indicates the location of the soma. **G:** A maximum projection image of an AAV2-GFP labeled Campana cell (green, G1) co-labeled with anti-Otx2 antibody (white, G1, and G2) shows that the Campana cell is Otx2-negative (2G2). The arrow indicates the location of the soma. **H:** A maximum projection image of an AAV2-GFP labeled Campana cell (green, H1, and H2) co-labeled with anti-PKC antibody (teal, H1-H3). A single optical section (H2 and H3) shows that the Campana cell (H2) is PKC-negative (H3). The arrow indicates the location of the soma. **I:** A maximum projection image of an AAV2-GFP labeled Campana cell (green, I1, and I3) co-labeled with anti-CaBP5 antibody (purple, I1-I3). A single optical section (I2 and I3) shows that the Campana cell is CaBP5-negative (I2). The arrow indicates the location of the soma. See also supplemental video 1-3 associated with this figure.

One feature common to BCs is that all BCs use glutamate as their neurotransmitter and transport glutamate from the cytosol into pre-synaptic vesicles in their axons via vesicular glutamate transport 1 (VGluT1) (Sherry et al., 2003; Johnson et al., 2004). Another feature common to BCs is that they express ribbons at their axonal terminals to regulate glutamate release (West, 1976; Vogel et al., 1977; Baden et al., 2013b). Accordingly, we labeled VGluT1 and ribbons in the wild type mouse retina with anti-VGluT1 (Sherry et al., 2003; Johnson et al., 2004) and anti-CtBP2 antibodies (Schmitz et al., 2000) to determine if Campana cells share this trait with BCs. The results show that both VGluT1 and CtBP2 are expressed by Campana cells in both the ON and OFF sublamina (Figures 2C and 2D) and that VGluT1 and CtBP2 are colocalized inside the Campana cell’s neurites in the IPL (Figures 2D). Therefore, we will refer to the Campana Cell’s neurites in the IPL as axons and predict that Campana cells are likely to release glutamate through ribbons synapses at their axonal terminals.

Next, we quantified the number of ribbon synapses within the Campana cell using anti-CtBP2 and 3D colocalization. Unlike other BCs, which primarily express synaptic proteins either in the ON or OFF layer of the IPL, the Campana cells have a similar number of ribbon synapses in the ON and OFF sublaminae of the IPL (Figure 2E and Figure 2— supplemental video 2). Although, the total number of ribbon synapses (27.8 ± 4.1) is on the low end when compared to other BCs, which range from 31 ribbon synapses (type 9) to 170 ribbon synapses (type 7) (Tsukamoto and Omi, 2017). Therefore, Campana cells might release glutamate into both the ON and OFF pathways, which is in contrast to other BCs that predominately form synapses in either the ON or OFF sublamina (Euler et al., 2014; Tsukamoto and Omi, 2017).

In addition, we tested a few common BC antigens to determine if the Campana cell expresses any of these, and the results show that Campana cells do not express two pan-BC antigens, Vsx2, and Otx2 (Figures 2F and 2G). Nor do they express the rod BC antigen, PKC (Figure 2H), or the cone BC antigen, CaBP5 (Figure 2I).

### mGluR6 is functional in the Campana cell

Since the Campana cells appear to express the mGluR6 at their dendrites postsynaptic to photoreceptors, we wanted to determine if this receptor functions to detect glutamate release from photoreceptors. Accordingly, we generated an AAV2-CAG-GCaMP6m (AAV2-GCaMP6m) viral vector (Figure 3A-3C) to measure the light-evoked changes in intracellular calcium of Campana cells using a two-photon imaging system.

**Figure 3.**
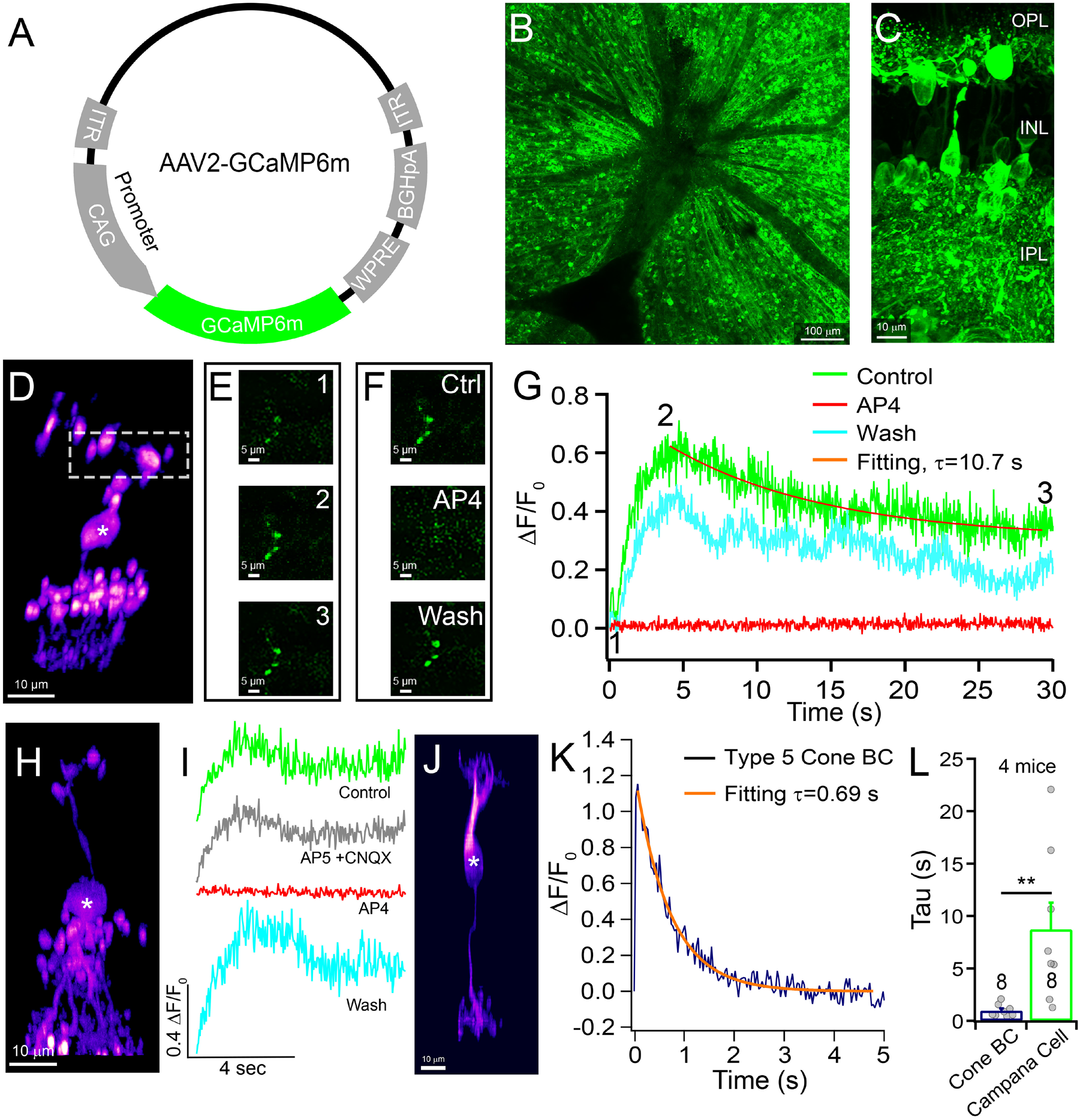
mGluR6 is functional in the Campana cell. A: Structure of the AAV2-CAG-GCaMP6m viral cassette. **B:** A view of an area of a flat-mount retina treated with the AAV2-CAG-GCaMP6m (AAV2-GCaMP6m) viral vector shows wide expression of GCaMP6m across the retina. **C:** A retina section from a mouse treated with AAV2-GCaMP6m. **D:** An AAV2-GCaMP6m expressing Campana cell. Its light responses were recorded at its dendrites (dashed box) and are shown in 4E-4G. **E:** A fluorescent image of the Campana cell dendrites at the initial image (E1, point 1 in 4G), at the peak intensity (E2, point 2 in 4G), and 30 seconds into the acquisition (E3, point 3 in 4G). **F:** The peak fluorescent intensity of the Campana cell before (Ctrl), during bath application of AP4, and 20 minutes after washing out AP4 (wash). **G:** ΔF/F0 measured at the dendritic terminal of the Campana cell in control (green), during bath application of AP4 (red), and 20 minutes after the washout of AP4 (teal). An exponential decay fitting (orange) was used to measure τ. The brief bump in the green trace is a light-induced artifact. **H:** An AAV2-GCaMP6m expressing Campana cell. Its light responses were recorded at its soma. **I:** Change in the light-evoked fluorescent intensity from the Campana cell in 4H during control (green), bath application of AP5 and CNQX (grey), bath application of AP4 (red), and 20 minutes after the washout of AP4 (teal). **J:** An AAV2-GCaMP6m expressing type 5 cone BC. **K:** ΔF/F0 of the type 5 cone BC after light stimulation (purple). An exponential decay fitting (orange) was used to measure τ. **L:** A comparison of the mean τ values of type 5 cone BCs (0.98 ± 0.21 sec) and Campana cells (8.7 ± 2.6 sec). A student t-test showed a significantly higher τ for Campana cells (p = 0.0092). The number in each column indicates the number of cells. Data are represented as mean + SEM. Cells in 4D, 4H, and 4J were masked from other cells for clarity, and a gaussian filter was applied to the image. Somas are indicated by “*” in these panels. See also Figure S5. See also supplemental figure 1.

Figure 3D shows a GCaMP6m expressing Campana cell from a dark-adapted retina. The GCaMP6m activity at the dendritic terminals of this Campana cell in response to a UV light flash (dashed box, Figure 3D) is shown at the first scanning image immediately after the light offset (“initial;” Figure 3E1), at the peak intensity (Figure 3E2), and 30 seconds after the flash (Figure 3E3). ΔF/F0 (F0: initial fluorescent intensity; ΔF: the difference between the fluorescence intensity from each time point and F0) for the Campana cell was plotted as a function of time (Figure 3G, control) to show that this cell has an increased fluorescent signal following the light flash, which decays slowly over time, indicating a light-induced ON response (Figure 3—supplemental figure 1; see Methods for further details). Consistently, bath application of a mGluR6 receptor agonist, AP4 (2-amino-4-phosphonobutyrate) (Shiells et al., 1981; Slaughter and Miller, 1981) reversibly abolished the light-evoked response (Figures 3F and 3G). However, simultaneously blockade of NMDA, AMPA, and kainate receptors did not affect the light response (Figure 3H and 3I), demonstrating that this light response is evoked by direct synaptic input from photoreceptors. Surprisingly, the kinetics of the Campana cells’ light responses seem to be much slower than that of type 5 cone BCs (Figures 3J-3K). Type 5 cone BCs have a sharp increase in GCaMP6m fluorescence immediately after a light flash and return to baseline in less than 3 seconds (Figure 3K). However, the average decay time constant (τ) of Campana cells (8.7 ± 2.6 sec) is significantly slower than that of type 5 cone BCs (0.98 ± 0.21 sec, p = 0.0092, student t-test; Figure 3L).

### Campana cells have close synaptic contact with both rods and cones

We further determined if Campana cells synapse with photoreceptors as BCs do. BCs are divided into rod BCs and cone BCs based on whether they primarily receive synaptic inputs from rods or cones (Ghosh et al., 2004; Wässle et al., 2009; Euler et al., 2014; Behrens et al., 2016; Tsukamoto and Omi, 2017). We examined whether the synaptic connections of Campana cells are dominated by rods or cones by examining the proximity of the Campana cell dendrites to the synaptic ribbons of photoreceptors. We transfected Campana cells with the AAV2-GFP vector and co-labeled the retina with either anti-REEP6 (rods) or anti-cone arrestin (cones) antibodies (Liu et al., 2012; Hao et al., 2014; Veleri et al., 2017). Using a previously published technique for identifying synaptic contacts (Strettoi et al., 2018), we identified the rod or cone synapses that contacted the Campana cell dendrite. Our results showed that the dendrites of Campana cells are in proximity (≤300 nm away) with both rod (Figure 4A-4E) and cone synapses (Figure 4F-4I). For measuring the number of close contacts with cones and rods, we only used cells that had their entire dendritic structure included in a single retinal section (40 µm in thickness). Figure 4D shows a 90-degree rotation of the entire dendritic plexus from the dash-lined box in Figure 4C. We confirmed that each Campana cell that we analyzed was a Campana cell by the morphology of its axons and dendrite, with each being a single cell (Figure 4C and 4H). Statistically, the mean number of contacts between Campana cells and cones (5 ± 0.8, n = 8) is not different from that of rods (3.8 ± 0.98, n = 6, p = 0.38, unpaired student t-test; Figure 4I), demonstrating that Campana cells likely receive equal input from both rods and cones.

**Figure 4.**
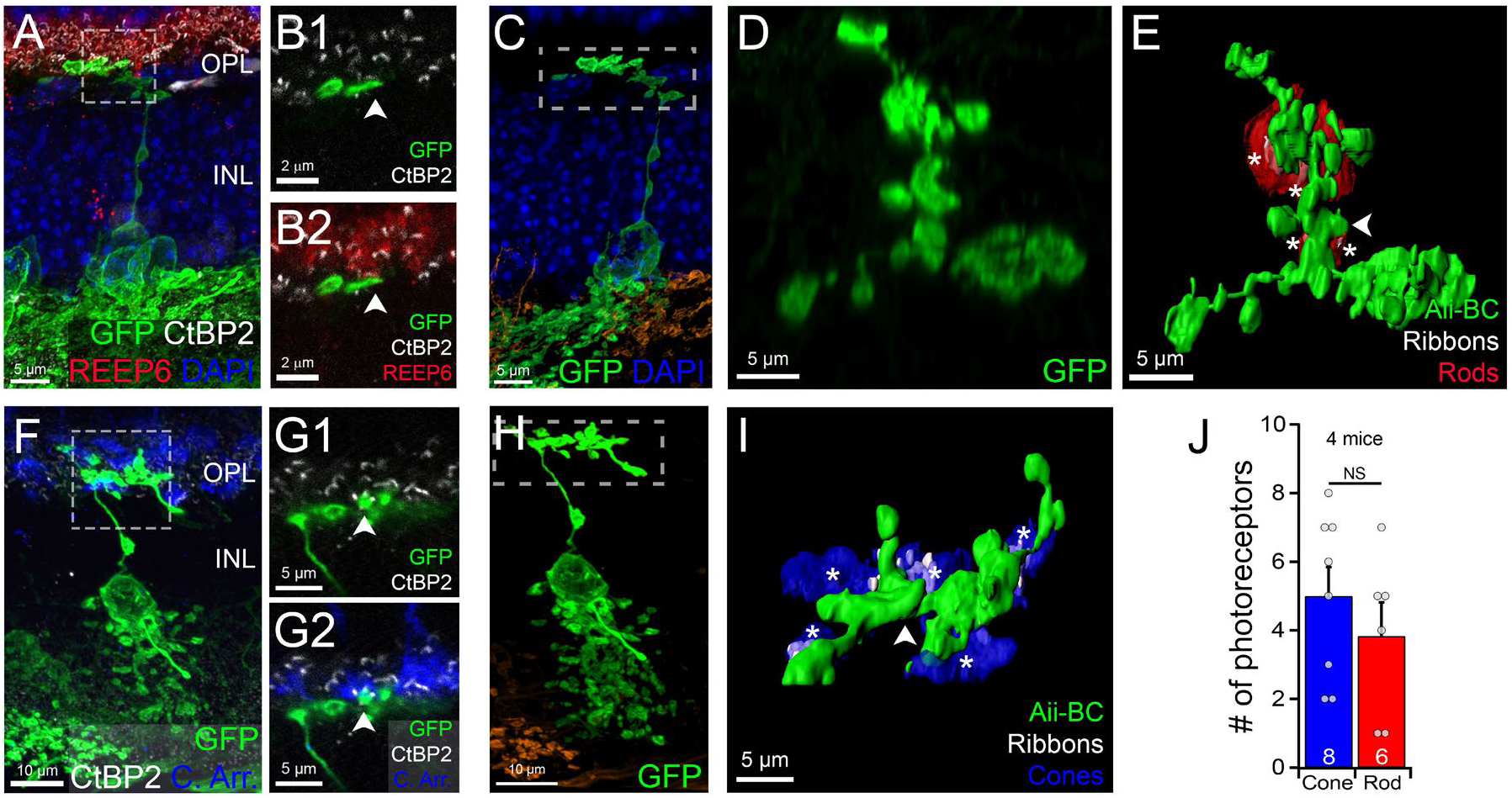
Campana cells have close synaptic contact with both rods and cones. A: An AAV2-GFP labeled Campana cell (green) co-labeled with anti-REEP6 (rods, red) and anti-CtBP2 (white) antibodies. **B:** A high-resolution single optic section of the boxed area in 5A shows that the dendrite of the Campana cell (green, B1, and B2) is in close contact with a ribbon synapse (white, B1 and B2) in a rod (red, B2). **C:** A masked view of the same Campana cell (green) shown in 5A confirms that it is a Campana cell. **D:** The entire dendritic plexus of the Campana cell in panel 5A and 5C with a 90° rotation along the X-axis. **E:** A surface rendering of the dendrite of the Campana cell as shown in panel D and rod photoreceptor terminals (red) with ribbon synapses inside (white) that are in close contact with the Campana cell dendrite. Asterisks indicate individual rod terminals, and the arrow indicates the dendritic segment marked with arrows in B1 and B2. **F:** An AAV2-GFP labeled Campana cell (green) co-labeled with anti-cone arrestin (cones, blue) and anti-CtBP2 (white) antibodies. **G:** A high-resolution single optic section of the boxed area in 5F shows that the dendrite of the Campana cell (green, G1, and G2) is in close contact with a ribbon synapse (white, G1 and G2) in a cone (blue, G2). **H:** A masked view of the same Campana cell (green) shown in 5F confirms that it is a Campana cell. **I:** A surface rendering of the dendrite boxed in 5H shows cone photoreceptor terminals (blue) with ribbon synapses inside (white) that are in close contact with the Campana cell dendrite (green). Asterisks indicate individual cone terminals, and the arrow indicates the dendritic segment of the Campana cell marked with arrows in G1 and G2. **J:** A comparison of the number of contacts to Campana cell dendrites from cones and rods. The cone contacts 5 ± 0.85 cones/Campana cell is not significantly different from that of rods (3.8 ± 0.98 rods/Campana cell, mean ± SEM, p = 0.38, unpaired student t-test). Dots represent individual cells, and numbers in each column are the number of cells.

### Campana cells receive functional inputs from both rods and cones

We then sought to quantify the synaptic inputs from rod and cone input to Campana cells. We recorded the intensity-response curve of Campana cells to light stimuli and compared their responses to those of type 5 cone BCs transfected with the AAV2-GCaMP6m virus. Figure 5A shows the morphology of a GCaMP6m^+^ Campana cell, and Figure 5B shows the light-evoked ΔF/F0 in GCaMP6m from the same Campana cell in response to both a bright UV light flash for cone responses and a dim green light flash for rod responses (see Figure 5—supplement figure 1). The Campana cell had responses with small variation in amplitude to all light intensities across three orders of magnitude of light intensities (Figure 5C). However, a GCaMP6m^+^ type 5 cone BC (Figure 5D) had no response to the lowest light intensity (Figure 5E), which is consistent with the notion that the green light stimulus at the intensity of 0.549 × 10^3^ photons/(μm^2^ × sec) does not activate cones (Figure 5F; n = 8). Indeed, the average max ΔF/F0 of Campana cells was 14.5 times higher than type 5 cone BCs (0.29 ± 0.04, Campana cell, n = 6, vs 0.02 ± 0.02, type 5, n = 8; unpaired student t-test, p < 0.0001), demonstrating that they have a high sensitivity to light. The response of type 5 cone BCs quickly increased with increasing light intensity before becoming fully saturated (Figure 5F), consistent with being a cone bipolar cell. Campana cells, on the other hand, showed consistent responses to both rod and cone-mediated inputs, corroborating what was seen through confocal imaging, that they receive synaptic inputs from both rods and cones.

**Figure 5.**
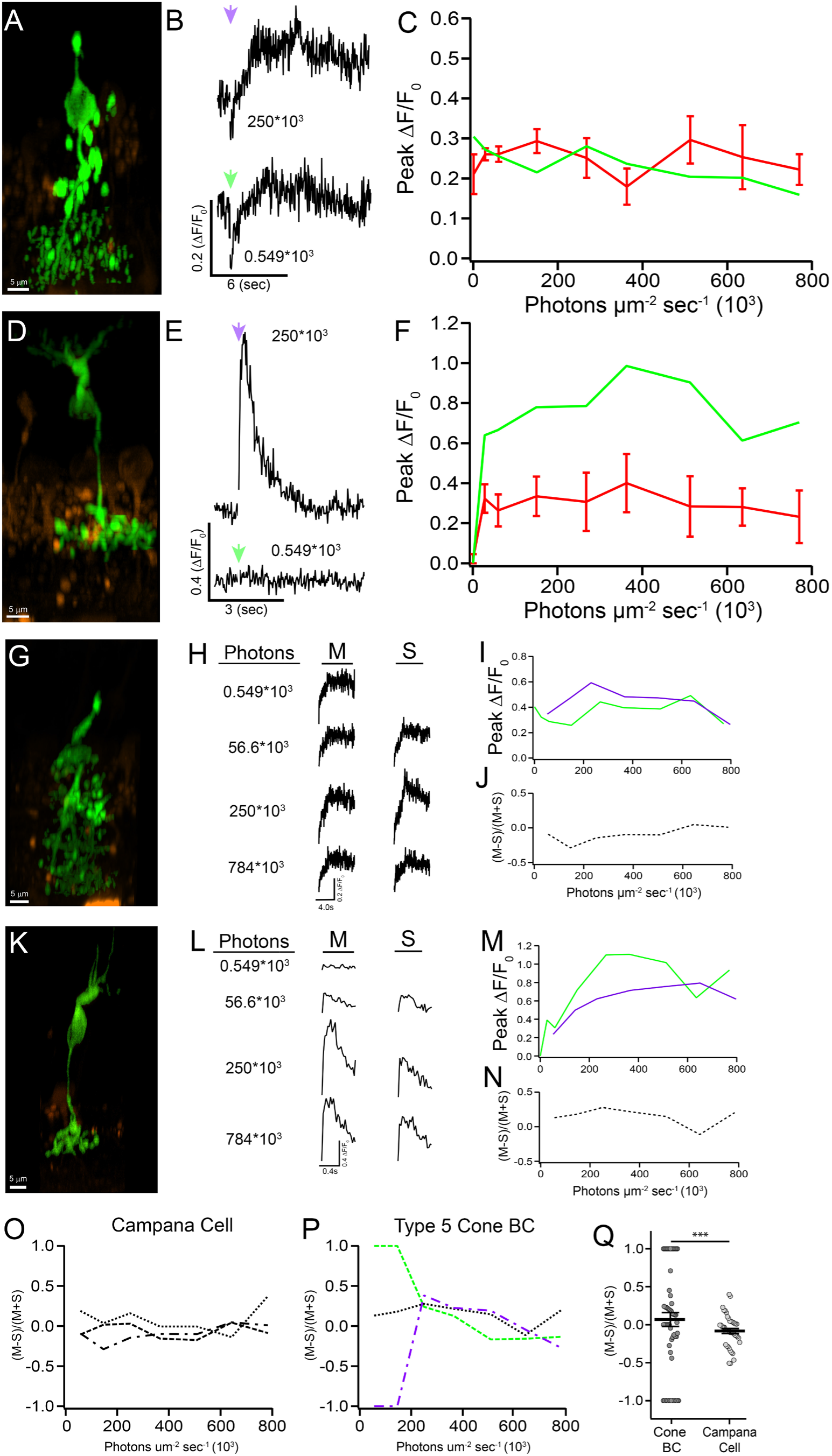
Campana cells receive functional inputs from both rods and cones. A: A Campana cell expressing GCaMP6m that had light responses recorded from its soma following the presentation of a 10ms light stimulus. **B:** The ΔF/F0 from the Campana cell in 5A evoked by a bright UV light stimulus (top, purple arrow), and a dim green light stimulus (bottom, green arrow). The early decrease in calcium ΔF/F0 is not due to inhibition but is due to a brief absence in two-photon laser scanning that results in a lower baseline (see methods and Figure 4—supplement figure 1:). **C:** The peak ΔF/F0 in response to 9 green light intensities of the Campana cell shown in panel 5A (green), and an average of 6 Campana cells (red) at each intensity. Photon measurements are in photons/(µm^2^ per second). **D:** A type 5 cone BC that had light responses recorded from its soma following a 10ms green light stimulus. **E:** The ΔF/F0 from the type 5 cone BC in 5D evoked by a bright UV light stimulus (top, purple arrow), and a dim green light stimulus (bottom, green arrow). **F:** The peak ΔF/F0 in response to 9 light intensities from the type 5 cone BC shown in 5D (green), and an average of 8 type 5 cone BCs (red) at each intensity. **G:** An Campana cell expressing GCaMP6m that had light responses recorded from its soma following the presentation of a 10ms light stimulus as shown in panels H-J **H:** The light-evoked ΔF/F0 of the Campana cell shown in panel G in response to green light stimuli at 4 intensities (M) and UV light stimuli at 3 intensities (S). Photon measurements are in photons/(µm2 per second). **I:** The peak ΔF/F0 of light-evoked GCaMP6m fluorescence signals in response to 9 green light intensities (green) and 7 UV light intensities (purple) of the Campana cell shown in panel G. **J:** A measurement of the response ratio of the green light stimuli (M) and the UV light stimuli (S) of the Campana cell shown in panel G. **K**: A type 5 cone BC expressing GCaMP6m that had light responses recorded from its soma following the presentation of a 10ms light stimulus, as shown in panels L-N. **L:** The light-evoked ΔF/F0 of the type 5 cone BC shown in panel K in response to green light stimuli at 4 intensities (M) and UV light stimuli at 3 intensities (S). Photon measurements are in photons/(µm2 per second). **M:** The peak ΔF/F0 of light-evoked GCaMP6m fluorescence signals in response to 9 green light intensities (green) and 7 UV light intensities (purple) of the type 5 cone BC shown in panel K. **N:** A measurement of the response ratio between the green light stimuli (M) and the UV light stimuli (S) of the type 5 cone BC shown in panel K. **O:** The response ratio of the green light stimuli (M) and the UV light stimuli (S) of 3 different Campana cells. All of them received comparative amount of synaptic inputs from both M- and S-opsins expressing cones. **P:** The response ratio of the green light stimuli (M) and the UV light stimuli (S) as a function of light intensity of 3 different type 5 cone BCs. One cell predominantly responds to the green light at low light intensity (green), one cell predominantly responds to UV light at low light intensity (purple), and the third cell received had a similar M- and S-opsins expressing cone input (black). **Q:** The distribution and average value of the response ratio to the green light stimuli (M) and the UV light stimuli (S) at 7 light intensities of type 5 cone BCs (8 cells) and Campana cells (6 cells). The variation in the light response ratio of Campana cells is significantly less than that of type 5 cone BCs (analysis of variance, F-test probability < 0.0001). Cells in 5A, 5D, 5G, and 5K were masked, and the background was colored in orange for clarity. Data are represented as mean + SEM in 5C, 5F and ± SEM in 5Q. See also supplemental figure 1 associated with this figure.

In addition, we examined whether Campana cells receive synaptic inputs from cones preferentially expressing M- or S-opsins. In pigmented C57/Bl6 mice, 34% of cones only express M-opsin, 26% of cones only express S-opsin, and 40% of cones express both M-opsin and S-opsin (Ortín-Martínez et al., 2014). To determine the inputs from cones preferentially expressing M- or S-opsin to Campana cells, we recorded light responses of Campana cells and type 5 cone BCs evoked by UV or green light stimuli. In these experiments, both Campana cells and type 5 cone BCs were randomly selected throughout the retina. To measure the relative strength of M-opsin and S-opsin inputs, we calculated the M/S input index based on the peak light responses to green and UV light stimuli using the following equation:

M/S input index = [Mi - Si] / [Mi + Si]

In this equation, M and S indicate the peak amplitudes of the cells to green and UV light stimuli at the light intensity (i). The M/S input index will be 1 for a cell only receiving M-cone inputs, it will be −1 for a cell only receiving S-cone inputs, and it will be 0 for a cell receiving an equal amount of input from M-opsin and S-opsin expressing cones. A Campana cell shown in Figure 5G responded to both green and UV light stimuli (Figures 5H and 5I) with an M/S input index that has a slight preference for UV at lower light intensities over the course of 7 different light intensities (Figure 5J). Figure 5K shows a side view of a GCaMP6m fluorescent image of a type 5 cone BC. As previously demonstrated, this cell did not respond to the green light stimulus at the intensity of 0.549 10^3^ photons/μm^2^ × sec (Figure 5L, top). However, with an increase in the light intensity, both the UV and green light stimuli elicited responses (Figures 5L and 5M). Figure 5N plots the M/S input index of this type 5 cone BC. Similarly, the M/S input index is roughly flat near 0, indicating similar amount inputs from both M and S-opsin expressing cones.

All Campana cells (n = 6) receive a similar amount of input from cones expressing M- and S-opsins and have similar M/S input index patterns as 3 different Campana cell M/S input index traces shown in Figure 5O. However, when we examined a group of type 5 cone BCs (n = 8), we found that type 5 cone BCs have a wide variety of M/S input index patterns, particularly at low light intensities. Figure 5P shows representative M/S input index plots of 3 type 5 cone BCs, in which one cell only receives M-opsin based responses at a low light intensity, another cell shows only S-opsin response under similar UV light intensities, and the third cell receives an equal input across all intensity levels. Statistically, analysis of the variation of M/S input index of type 5 cone BCs and Campana cells showed that type 5 cone BCs have a significantly higher variation in their M/S input index than that of Campana cells (Figure 5K; p < 0.0001, ANOVA). These results are consistent with previous reports that type 5 cone BCs are made up of a variety of BC subtypes that are difficult to distinguish morphologically but vary in genetic expression, photoreceptor input and light responses (Behrens et al., 2016; Shekhar et al., 2016; Franke et al., 2017; Tsukamoto and Omi, 2017).

### Campana cells express genes and proteins commonly expressed by ACs

Because Campana cells showed substantial morphological similarity to Aii-ACs but did not express pan-bipolar markers, we further examined if Campana cells express proteins generally expressed by ACs, especially by Aii-ACs. Our results show that Campana cells are positive for Pax6 (Figure 6A), a paired homeobox gene expressed in ACs, horizontal cells, and RGCs in the mouse retina (Hill et al., 1991; Belecky-Adams et al., 1997; de Melo et al., 2003; Bandah et al., 2007). Additionally, Campana cells are positive for calretinin (Figure 6B), a calcium-binding protein that is primarily expressed in ACs (Rogers, 1987; Völgyi et al., 1997; Gábriel and Witkovsky, 1998; Massey and Mills, 1999; Dyer and Cepko, 2001; Rösch et al., 2014; Lee et al., 2016), and glycine transporter 1 (GlyT1, Figure 6C and Figure 6—supplemental video 1), a protein that transports glycine from the extracellular space into cells, and is a known marker for Aii-ACs (Haverkamp and Wässle, 2000; Akopian et al., 2016; Pérez de Sevilla Müller et al., 2017).

**Figure 6.**
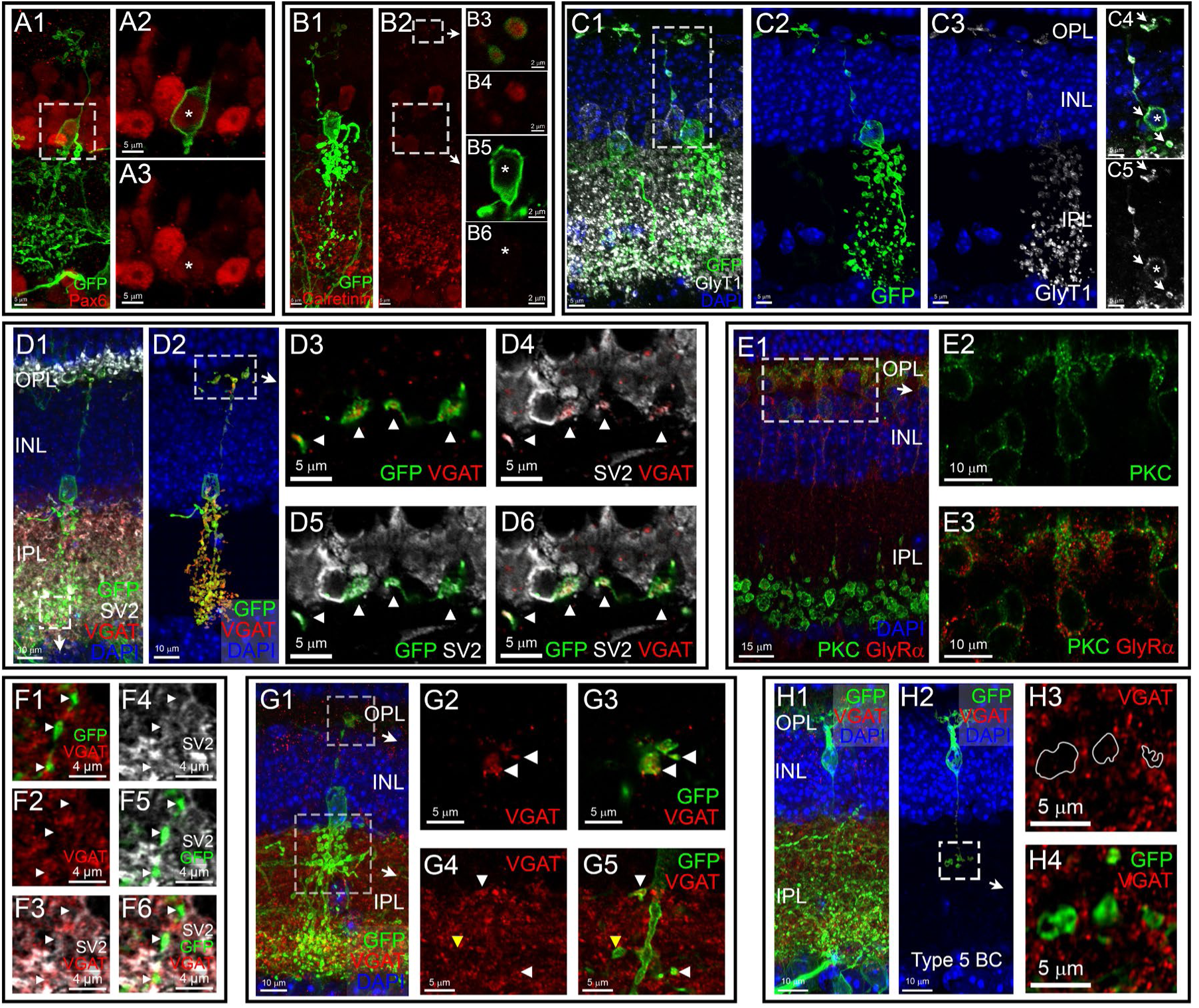
Campana cells express proteins commonly expressed by ACs. A: An AAV2-GFP transfected Campana cell (green) co-labeled with anti-Pax6 antibody (red). A single frame from the boxed area in A1 shows that the soma (asterisk) of the Campana cell is Pax6-positive (A2 and A3). **B:** A Campana cell (green) labeled by AAV2-GFP and co-labeled with anti-calretinin antibody (red, B1, and B2). A single frame from the boxed areas in B2 shows that the dendrites of Campana cells are calretinin-positive (red, B3 and B4), but the soma (asterisk) is not (B5 and B6). **C:** An AAV2-GFP transfected Campana cell (green cell, right side of C1) and Aii-AC (green cell, left side of C1) colabeled with anti-GlyT1 antibody (white, C1, and C3-C5). Masking of the Campana cell is shown in C2, and a masking of the anti-GlyT1 staining within the Campana cell is shown in C3. A single optical frame from the boxed area in C1 shows that the dendrites, soma (asterisk), and axons (arrows) are GlyT1-positive (white, C4 and C5). **D:** A Campana cell (green, D1) labeled by AAV2-GFP and co-labeled with anti-VGAT (red), and anti-SV2 (white; two adjacent sections were used for max projection) antibodies. A masked view of the same cell in D1 with GFP and anti-VGAT labeling (D2). A single super-resolution optic section of the area boxed in D2 shows that Campana cell dendrites are positive for SV2 and VGAT (D3-D6). D3: GFP and anti-VGAT staining. D4: Anti-SV2 and anti-VGAT staining. D5: GFP and anti-SV2 staining. D6: GFP, anti-SV2, and anti-VGAT staining. Arrowheads indicate the Campana cell’s dendrite. **E:** Mouse retina co-labeled with anti-PKC (green, rod BCs) and anti-GlyRα1+2 (red) antibodies shows that GlyRα overlaps with rod BCs (E1). Super-resolution imaging confirms that the dendrites of rod BCs (E2) express GlyRα (E3). **F:** A single super-resolution optic section of the area boxed in D1 shows that Campana cell axons are positive for SV2 and VGAT (F1-F6). F1: GFP and anti-VGAT staining. F2: Anti-VGAT staining. F3: Anti-SV2 and anti-VGAT staining. F4: Anti-SV2 staining. F5: GFP and anti-SV2 staining. F6: GFP, anti-SV2, and anti-VGAT staining. Arrowheads indicate the location of the Campana cell’s axon. **G:** A Campana cell (green, G1) labeled by AAV2-GFP and co-labeled with anti-VGAT antibody (red). A single super-resolution optic section of the area boxed in G1 (top) shows that the Campana cell’s dendrites are positive for VGAT (G2 and G3). A single super-resolution optic section of the area boxed in G1 (bottom) shows that VGAT (G4 and G5) is located inside some (white arrow heads), but not all (yellow arrow head), of the Campana cell’s axons. **H:** A GFP-positive type 5 cone BC (green, H1) co-labeled with an anti-VGAT antibody (red, H1-H4). A masked view of the same cell shows no VGAT protein in the cell (H2). A single super-resolution optic section from the boxed area in H2 shows that the axons (green/outline) of the type 5 cone BC are negative for VGAT (red, H3-H4). See also supplemental video 1 associated with this figure.

BCs are excitatory neurons and only release glutamate as their neurotransmitter. However, Campana cells express GlyT1, which would enable Campana cells to transport glycine from the extracellular space into their cytosol. To further determine if Campana cells could release glycine as a neurotransmitter, we examined whether Campana cells express the vesicular inhibitory amino acid transporter (VIAAT/VGAT), which is a membrane transport protein that transports both glycine and GABA from the cytosol into synaptic vesicles (McIntire et al., 1997). When we co-labeled AAV2-GFP transfected GFP^+^ Campana cells with anti-VGAT and anti-SV2 antibodies (McIntire et al., 1997; Wan et al., 2010 p.2; Weltzien et al., 2012), we found that every GFP^+^ Campana cell was VGAT positive and VGAT colocalized with synaptic vesicles labeled by anti-SV2 antibody (Figure 6D). VGAT expression was primarily in the dendritic and axonal terminals of Campana cells (Figures 6D2-6D6) but not in other BCs (data not shown). Since antibody staining showed that Campana cells do not express glutamate decarboxylase (GAD, data not shown), which synthesizes GABA (Lakowski et al., 2007; Walls et al., 2010), we concluded that VGAT packages synaptic vesicles in Campana cells with glycine. While Campana cells likely release glycine in the OPL, and the presence of GlyRα in the OPL has been previously shown (Heinze et al., 2007), we determined what cells express glycine receptor α (GlyRα) in the OPL. We found that GlyRα was localized primarily in the dendrites of rod BCs in the OPL, as shown by the colocalization of anti-PKC and anti-GlyRα1+2 staining (Figure 6E). Therefore, Campana cells could inhibit rod BCs through activation of their dendritic glycine receptors. Additionally, Campana cells are VGAT positive in their axons as well (Figure 6F and 6G). However, VGAT is punctate and does not fill every axon, but SV2 does seem to (Figure 6F). We also found that as is to be expected, other BCs, such as a type 5 cone BC (Figure 7H) are negative for VGAT, and confirmed this with super resolution imaging (Figure 6H3 and 6H4). This indicates that Campana cells are unique in the retina for having vesicular transporters for glycine as well as glutamate.

**Figure 7.**
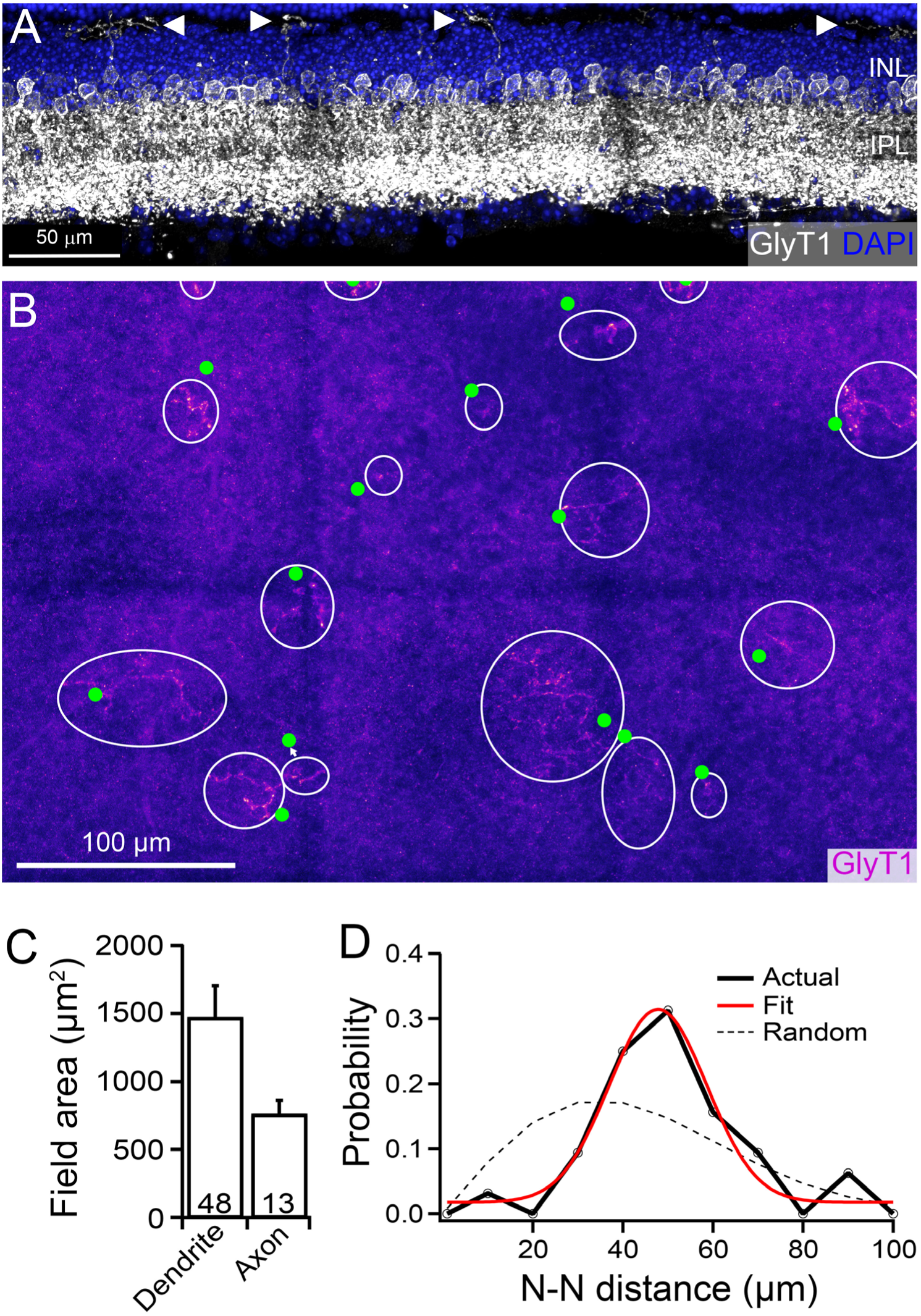
Campana cells have a sparse, non-random retinal mosaic. A: A mouse retina cross-section labeled with anti-GlyT1 antibody (white) and DAPI (blue) shows the dendrites of 4 different Campana cells (arrowheads) projecting to the OPL. **B:** A view of the OPL of a mouse retina flat-mount shows the dendrites of several Campana cells. The dendritic field area is outlined with white circles, and the approximate location of each soma is indicated by green circles. **C:** The average field area of the dendrites and axons of Campana cells. The dendritic field area measurements were taken from Campana cells in flat-mount retinas that were labeled with anti-GlyT1 antibody or AAV2-GFP (5 mice). Axonal field area measurements were taken from Campana cells in flat-mount retinas that were labeled with AAV2-GFP (4 mice). The numbers in each bar are the number of cells. Data are represented as mean + SEM. **D:** A probability distribution function for the Nearest Neighbor distance of each Campana cell soma. The solid black line is the actual observations in bin widths of 10 µm, the solid red line is a normal gaussian distribution, and the dashed line is a random simulation based on cell density. Mean = 55.2 ± 2.91 µm, Conformity ratio/Regularity index = 3.35, n = 32, 2 mice.

### Campana cells have a sparse retinal mosaic

Since every AAV2-GFP transduced Campana cell expresses GlyT1 throughout the cell (Figure 6C), and Campana cells are the only GlyT1^+^ cell type that has dendrites projecting into the OPL (Figure 7A), we used anti-GlyT1 labeling to estimate the density of Campana cells in a flat-mount retina of adult mice (Figure 7B). We found that Campana cells have a density of 132 ± 5 cells/mm^2^ (mean ± SEM, 953 Campana cells, 4 mice).

We next measured the dendritic and axonal field areas of the Campana cells to determine the coverage area on the retina (Figure 7C). We obtained dendritic field area measurements from Campana cells in flat-mount retinas from mice that were labeled with anti-GlyT1 or AAV2-GFP and the axonal field area from Campana cells in flat-mount retinas that were labeled with AAV2-GFP. The average dendritic field area was 1469 ± 233.7 µm^2^ (n = 48, 5 mice) and the average axonal field area was measured as 757.2 ± 105.5 µm^2^ (n = 13, 4 mice). While the dendritic field size has a large variance (range = 65.0 – 6457 um^2^ as seen in Figure 7B), the axonal field size is much more consistent (range = 409.0 – 1635 um^2^). On average the Campana cell’s dendritic field size is ∼10 times larger than that of rod BCs (Keeley and Reese, 2010), ∼7 times larger than type 5 cone BCs (222 ± 35.5 μm^2^, n = 9, 5 mice), ∼12 times larger than type 6 cone BCs (Dunn and Wong, 2012), ∼7 times larger than type 7 cone BCs (Dunn and Wong, 2012), and ∼2 times more massive than type 8 cone BCs (Dunn and Wong, 2012). Indicating that Campana cells likely have a much larger dendritic field area than ON BCs. It has been postulated that each of the retina’s neuronal types is regularly spaced so that each of the cell types covers the retinal surface evenly to survey the visual scene efficiently (Wässle and Riemann, 1978; Wässle et al., 1981; Cook, 1996; Masland, 2012). This coverage can be measured by the coverage factor, determined by the spatial density of the cells (cells/mm^2^) times the dendritic field area of each cell (mm^2^/cell). We calculated the coverage factor for Campana cells as 0.46 for the dendrites and 0.24 for the axons. We estimated the coverage factor of type 5 cone BCs based on the density previously reported density measurements (Wässle et al., 2009) as 1.11. Additionally, type 7 cone BCs and rod BCs were estimated to have coverage factors of 0.78 and 2.65, respectively (Wässle et al., 2009; Keeley and Reese, 2010). Therefore, the Campana cell has a lower coverage factor than other BC types.

To determine if the Campana cells have regular spacing, we measured the nearest neighbor distance between somas labeled by GlyT1 (see Figure 7B green circles). The mean nearest-neighbor (N-N) distance is 55.2 ± 2.91 µm, and the median is 53.2 µm. The N-N probability distribution fits with a gaussian curve, but not a random distribution based on the cell density, indicating non-random spacing (n = 32, 2 mice; Figure 7D) (Wässle and Riemann, 1978). Additionally, to determine if the Campana cells have regular, non-random spacing we calculated the Conformity ratio/Regularity index (mean/SD) (Wässle and Riemann, 1978; Cook, 1996). The calculated conformity ratio of 3.35 indicates that Campana cells have a non-random distribution in the retina (p < 0.001) (Cook, 1996).

### Campana cells are genetically unique

Furthermore, we examined the RNA expression pattern of Campana cells using single-cell RNA seq analysis described previously (Daines et al., 2011; Zhao et al., 2016), and compared the results with Aii-ACs and rod BCs. Using a patch-clamp electrode, we collected 5-10 Campana cells, Aii-ACs transduced by AAV2-GFP (Figures 8A-8C), and rod BCs transduced by AAV2.7m8-mGluR6-mCherry (AAV2.7-mCherry; Figures 8D and 8E) viral vector (Lu et al., 2016). The RNAseq library was generated according to the SMART-seq v4 Ultra-low input RNA kit (Takara Clontech). In brief, isolated cells were lysed in lysis buffer, 3-SMART-seq CDS primer II and V4 oligonucleotide were added for first stranded cDNA synthesis. cDNA was amplified using PCR Primer II A, and subsequently purified using Ampure XP beads (Beckman). Illumina library was prepared using the Nextera XT DNA library preparation kit (Illumina) and sequenced using Illumina Novaseq (Daines et al., 2011; Zhao et al., 2016). The sequencing analysis was done blind to the cell type. We sequenced 34,118,309 reads from these cells and 20,289 actively expressed transcripts were detected. From these actively expressed transcripts, 6913 are uniquely expressed by the 3 types of neurons (6301 by Aii-ACs, 415 by Campana cells, and 197 by rod BCs). Next, we used FPKM (Fragments Per Kilobase of transcript per Million mapped reads) profile for a comparison between the samples after removing unexpressed transcripts across all the samples (13,237 transcripts) and performing quantile normalization (limma package, R). Pairwise Pearson correlation analysis shows that the expression patterns of Aii-ACs and Campana cells are positively correlated (r = 0.7). However, the gene expression patterns of Campana cells to rod BCs (r = 0.4) and Aii-ACs to rod BCs (r = 0.45) are not correlated (Figures 8F and 8G). Additionally, we looked at the transcripts that are shared or unique to each cell type. From this analysis, it was clear that both Campana cells and rod BCs share a majority of their transcripts with Aii-ACs (Figure 8H).

**Figure 8.**
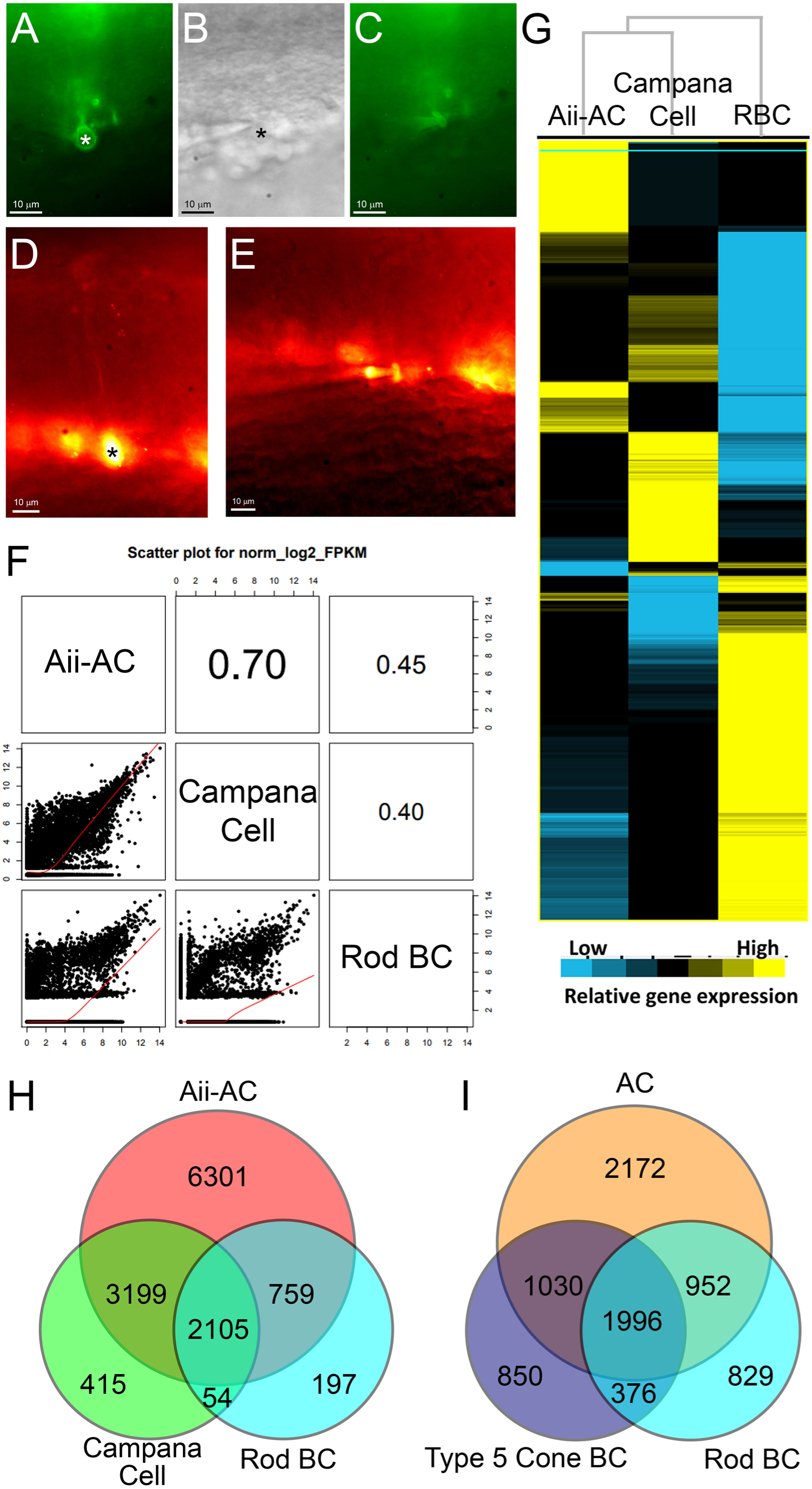
Campana cells are genetically unique. A: An Aii-AC (green) labeled with AAV2-GFP in a live retinal slice. An asterisk marks the soma. **B:** Same area in panel A in brightfield view. The cell collection needle is visible to the left. An asterisk marks soma. **C:** Same area in panels A and B after negative pressure was applied to the needle to collect the cell contents. **D:** A rod BC (red) labeled with the AAV-mGluR6-mCherry viral vector in a live retinal slice. **E:** The same cell marked in panel D after the cell contents were sucked into the needle. **F:** The FPKM profile was used for cell type comparison. Pairwise comparison across the cell types shows that Aii-ACs and Campana cells are more closely correlated (0.70) than either of them are to rod BCs (0.45 and 0.40, respectively). **G:** The hierarchical clusterings of the expression patterns of Aii-ACs, Campana cells, and rod BCs. Each column represents a cell type, and each row represents a gene. The scale at the bottom indicates the relative level of expression. **H:** A Venn Diagram showing the RNA reads that are shared by, and unique to each group. The number indicates the number of reads in each section. **I:** A Venn Diagram showing the RNA reads that are shared by, and unique to each group from a previously published and publicly available database (Shekhar et al., 2016). A random sample of 10 cells was taken from the amacrine cell cluster, type 5a cone BC cluster, and rod BC cluster. The number indicates the number of reads in each section.

We were, therefore, curious to address the question of how much does RNA expression vary between cells of different types or between neurons within the same class? To address this point, we analyzed a previously published database of BC RNA-seq (Shekhar et al., 2016). All the data from this roughly 27,000 cell sequencing experiment is available online (https://singlecell.broadinstitute.org/single_cell/study/SCP3/retinal-bipolar-neuron-drop-seq#study-summary). When we took a random sample of 10 cells from each BC cluster (140 cells, 14 clusters in total) and compared it with 10 cells from the AC cluster in the same data set. We found that all BC types had 70-78% of their RNA transcripts in common with the AC cluster, BCs however, only shared 56-63% of their transcripts with any other type of BC cluster (Figure 8I). Therefore, BCs share more transcripts with ACs than they do with each other. Consistently, Shekhar et al. (2016, figure 1C) shows that some BCs can cluster closer to ACs than they do to other BC types (type 2 for example) and that previously accepted distinct cell types by morphology were not significantly distinguishable by RNA reads alone (type 2 from 1a/b and types 8/9 from each other, figure 1G).

### Campana cells are evolutionarily conserved in mammals

Finally, we determined if Campana cells are a cell type preserved in other mammals by labeling the retinas of marmosets and macaques with anti-GlyT1 antibody. Figure 9A shows a macaque retina co-labeled by an anti-GlyT1 and an anti-CtBP2 antibody. Just as in the mouse, the macaque retina contains GlyT1^+^ cells with dendrites projecting to the OPL, and this cell also has an axonal plexus ramified in both ON and OFF sublaminae of the IPL as was seen in Campana cells in mice (Figure 9B). Additionally, the dendrites of the GlyT1^+^ cell appear to be in near proximity with the ribbon synapses of both rods and cones (Figure 9B). Similarly, co-labeling of marmoset retina using anti-GlyT1 and anti-CtBP2 antibodies revealed GlyT1^+^ cells with a morphology resembling what we observed in the macaque and mouse retina (Figure 9C).

**Figure 9.**
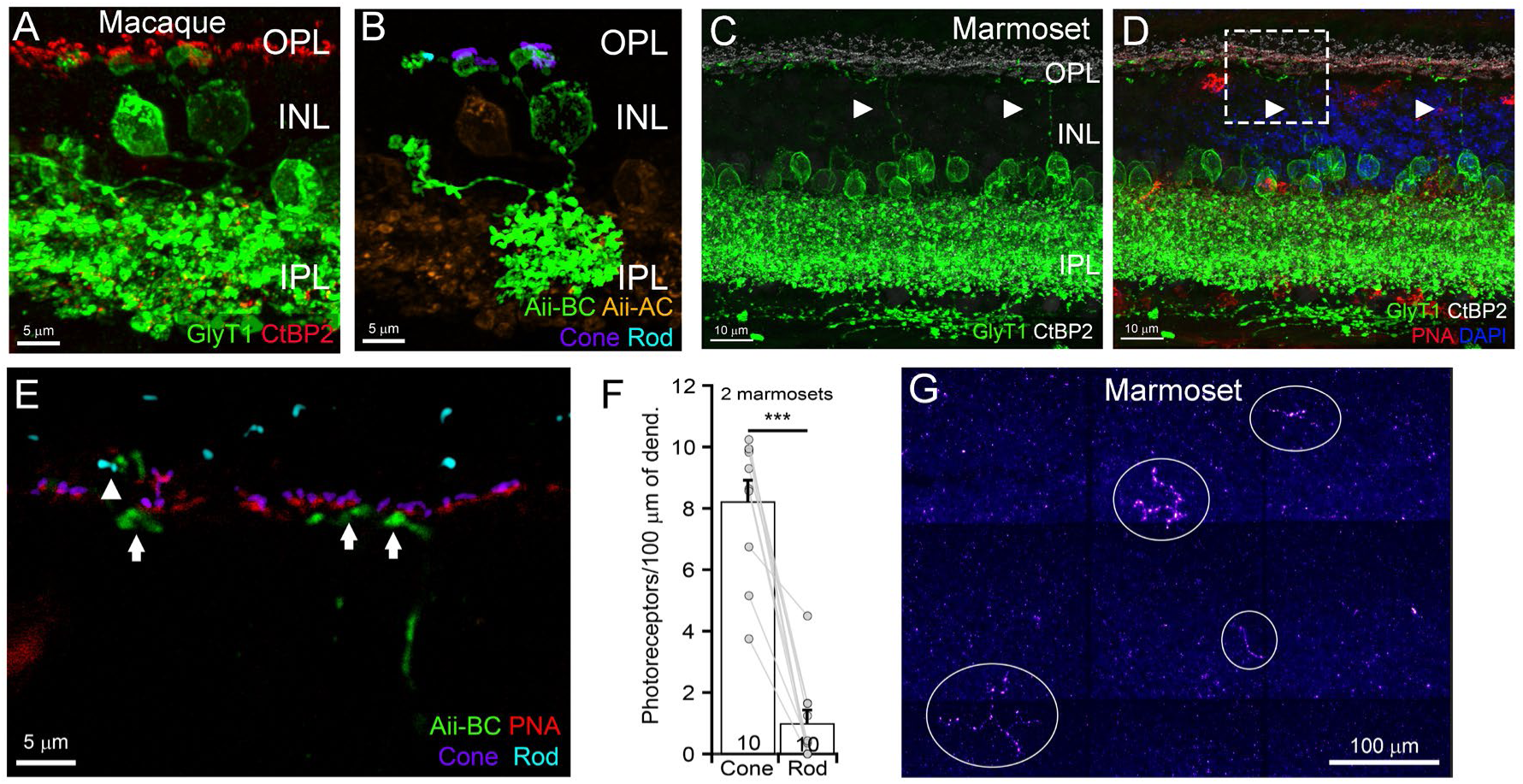
Campana cells are evolutionarily conserved in mammals. A: A macaque retina labeled with anti-GlyT1 (green) and anti-CtBP2 (red) antibodies. **B:** The same view in panel A, but with the Campana cell masked and isolated (green) from the Aii-ACs (orange). The dendrites show close contacts with 2 cone ribbons (purple) and 1-rod ribbon (teal). **C:** A marmoset retina co-labeled with anti-GlyT1 (green) and anti-CtBP2 (white) antibodies shows two Campana cells. **D:** The same view in panel C, but with DAPI (blue) and PNA-Alexa555 (red) to label cone terminals. **E:** A single frame view of the area boxed in D shows that the dendrites of the Campana cell have close contacts with both cone ribbons (purple, arrows) and a rod ribbon (teal, arrowhead). **F:** A comparison of the density of close contacts between the dendrites of Campana cells to rods or cones. A paired student t-test shows that the mean density of cone contacts (8.2 ± 0.7 cones/100 µm of dendrite) is significantly higher than rod contacts (0.98 ± 0.4 rods/100 µm of dendrite; p < 0.0001, paired student t-test). Dots represent individual cells, and numbers in each column are the number of cells. Data are represented by mean + SEM. **G:** A view of the OPL of a marmoset retina flat-mount shows the dendrites of several Campana cells. The dendritic field area is outlined with white circles.

We then used peanut agglutinin (PNA) fused to Alexa 555 to identify cone pedicles in the marmoset retina (Figure 9D) (Blanks and Johnson, 1984; Dkhissi-Benyahya et al., 2001). With this method, we characterized the close proximity between the dendrites of Campana-like cells (Figure 9E), and the rod and cone ribbon synapses. We found that in the primate retina there is a significantly higher density of synaptic contacts with cone ribbons than with rod ribbons (Figure 9F), likely due to the much lower rod density in the marmoset retina than the mouse retina (Jeon et al., 1998; Springer et al., 2011). Additionally, marmosets have a low density and wide variation in Campana-like cell dendritic field areas, just as in mice (Figure 9G). Nonetheless, our results demonstrate that Campana cells are preserved across species from mice to primates.

## Discussion

### What cell class do Campana cells belong to, BCs or ACs?

Is a Campana cell a single cell, or is it an amacrine cell fused with a bipolar cell? The Campana cell was found using a transsynaptic labeling technique that utilizes WGA as a critical component. It could be that this cell is a combination of a BC and an AC that are fused via a gap junction with WGA passing through to conduct labeling. However, WGA has never been reported to pass through a gap junction, and it is much too large to do so. Indeed, WGA exists as a homodimer with a measurement of 23.6 kDa in molecular mass (Finne, 1997), but gap junctions will only pass molecules with a molecular mass up to 1 kDa (Gong and Nicholson, 2001; Harris, 2001). Additionally, we were able to find the Campana cell in wild type mice as well as two primate species using the GlyT1 antibody, and in mice using the AAV2-CAG-ChR2-GFP-Na1.6 AAV2-CAG-GCaMP6m virusus. All of the Campana cells shown in this manuscript were labeled with these viruses or the GlyT1 antibody. All of this combined with the fact that every Campana cell soma was only connected to a single dendrite eliminates the possiblity that the Campana cell is actually two fused cells. Every Campana cell used for analysis in this work was viewed in three dimensions for verification.

A fundamental finding of this study revealed a previously undescribed unique type of retinal interneurons. This cell type shares some essential anatomical, physiological and biochemical features with BCs, including that the Campana cells have dendrites ramified in OPL, express mGluR6 at their dendritic terminals as other ON BCs (Nakajima et al., 1993; Nomura et al., 1994), and form synaptic contacts with photoreceptors. Campana cells also form direct contacts with inner retinal neurons through ribbon synapses and express VGluT1 near their ribbon synapses (Sherry et al., 2003; Johnson et al., 2004). Campana cells respond to a light stimulus with an increase in intracellular calcium, through direct photoreceptor input just as other ON cone BCs and rod BCs do (Shiells et al., 1981; Slaughter and Miller, 1981).

However, Campana cells also express several unique structural, functional, and genetic features, which are not present in any other BCs and do not follow several principles of the current paradigm of how BCs process visual signals (Masland, 2012). First, conventional BCs are divided into two major groups based on their synaptic inputs in all vertebrates, the rod BCs and cone BCs (Haverkamp et al., 2003; Euler et al., 2014). Under normal conditions, rod BCs predominately synapse with rods, while cone BCs primarily synapse with cones with rare exceptions (Kolb, 1970; Masland, 2012; Euler et al., 2014; Behrens et al., 2016; Santina et al., 2016; Tien and Kerschensteiner, 2018). In contrast, Campana cells synapse with a similar number of rods and cones in the mouse retina and do not seem to be selective for S- or M- cones. Even in a species with a low rod density, such as marmosets, Campana cells also appear to synapse with both rods and cones. Second, conventional BCs stratify their axonal terminals in either the ON or OFF sublamina in the IPL and selectively synapse with either ON or OFF ACs and RGCs (Masland, 2012). However, Campana cells ramify their axonal terminals throughout the entire IPL and form similar number of ribbon synapses with neurons in both the ON and OFF sublaminae. Third, all BCs are considered excitatory neurons and only release glutamate as their neurotransmitter (Sherry et al., 2003; Baden et al., 2013b, 2013a; Euler et al., 2014), while Campana cells express glycine transporters for both trans-membrane transporting and trans-vesicular transporting in addition to the expression of VGluT1. Accordingly, they are very likely to be the only retinal neurons that release both glutamate and glycine. Finally, the gene expression profile of Campana cells is significantly different from that of conventional BCs and they do not express several pan-BC markers, such as Vsx2 and Otx2, or BC type-specific markers, such as PKC or CaBP5. Therefore, Campana cells are significantly different from any BC type in many aspects although they primarily function as BCs do, to relay visual signals from photoreceptors to neurons in the inner retina.

Are Campana cells a type of AC? Although Campana cells express several AC specific markers, such as Pax6, Calretinin, GlyT1, and VGAT (Massey and Mills, 1999; de Melo et al., 2003; Lee et al., 2016; Remez et al., 2017; Bleckert et al., 2018; Eulenburg et al., 2018), and their axons resemble the morphological pattern of Aii-AC neurites, they are different from ACs in several fundamental aspects. First, Campana cells directly receive synaptic inputs from photoreceptors and relay visual signals to ACs and RGCs, and there is no AC that receives direct synaptic input from photoreceptors. Although some ACs, such as interplexiform ACs, project to the OPL from the INL and synapse with HCs in the OPL, they do not receive synaptic input from photoreceptors (Marc and Liu, 1984; Witkovsky et al., 2008; Dowling, 2012). Indeed, a cell with a similar morphology to the Campana cell has been previously observed at least twice in mice and described as an interplexiform cell (Fisher, 1979; Haverkamp and Wässle, 2000); however neither of these results looked at the cell’s functionality. Second, Campana cells express several BC specific synaptic proteins, such as CtBP2, mGluR6, and VGluT1, which are not expressed by any ACs. One might argue that since the gene expression pattern of Campana cells is positively correlated to Aii-ACs, they are genetically a type of ACs. However, our analysis of a previously published database of BC RNA-seq (Shekhar et al., 2016) demonstrated that BCs share more transcripts with ACs than they do with other BC types, and some BCs can cluster closer to ACs than they do to other BC types. Therefore, the correlation strength of gene expression between cell types might not be used as the sole source to distinguish cell types. Taken together, Campana cells share some fundamental features with both BCs and ACs but differ significantly from both in many other critical aspects. Therefore, Campana cells seem to not belong to either the BC or AC class.

Then, do Campana cells belong to a new retinal cell type? One might argue that Campana cells might be sporadic variations of conventional BCs or ACs. Because the Campana cells are preserved across species from mice to primates, they are unlikely to be the results of sporadic variations of conventional BCs or ACs in mice due to inbreeding or transgenic treatments. Interestingly, Pax6, a protein that Campana cells express, is required for the development of late-born retinal interneurons (including BCs and some ACs) (Bandah et al., 2007; Remez et al., 2017), and overexpression of Pax6 during retinal development leads to the development of glycinergic amacrine cells (Remez et al., 2017). This indicates that it is possible that Campana cells are formed during the wave of BC and glycinergic amacrine cell formation, but this possibility will require further investigation.

Although Campana cells have a low density, they are not unique in this point since several other neuronal types in the retina have a similarly low density including interplexiform ACs (∼29 cells/mm^2^, assuming a 2.5 mm radius mouse retina) (Sankaran et al., 2018), HCs (135-225 cells/mm^2^) (Wässle and Riemann, 1978), alpha RGCs (174 cells/mm^2^) (Wässle and Riemann, 1978), type 5d/X BC (∼385 cells/mm^2^) (Helmstaedter et al., 2013), and type 8 BC (∼333 cells/mm^2^) (Helmstaedter et al., 2013). Additionally, even with a low density, Campana cells have a non-random spatial distribution in the mouse retina. Additionally, the identification of the Campana cell in two separate primate species indicates that the Campana cell is not a random mutation in an inbred mouse strain. Therefore, Campana cells are likely to belong to their own unique retinal class.

### What role might Campana cells play in vision?

Another fundamental question is, what might be the functional role of the Campana cells in visual signal processing? Parallel processing is an organizing principle of many neural circuits. In the vertebrate retina, neuronal circuits process scotopic and photopic vision through rod- and cone-mediated synaptic pathways. Toward this end, rods and cones form synapses with distinct BC types, rod BCs or cone BCs (Mariani, 1984a, 1984b; Wässle and Boycott, 1991; Kolb et al., 1992). Accordingly, rod and cone BCs have distinguishable sensitivity to light intensity to process visual signals under scotopic and photopic conditions, respectively (Bloomfield and Völgyi, 2009; Dunn and Wong, 2014; Zele and Cao, 2015). However, Campana cells synapse with both rods and cones, which extends the range of their intensity-response relationship and results in a flat, non-monotonic intensity-response curve. It is possible that this rod/cone input could be a result of gap junction coupling through rods and cones (Bloomfield and Dacheux, 2001), however, the lack of a rod response from type 5 cone BCs under the same conditions serve to refute that this is what we observed. This dual rod/cone input enables the Campana cells to transmit light responses of both rods and cones to the inner retina but prevents Campana cells from distinguishing the synaptic input of rods and cones. Therefore, although Campana cells relay visual signals from photoreceptors to inner retinal neurons like other BC types, the signals transmitted by Campana cells will likely play different roles in visual signaling processing than other BC types. In addition, the slow kinetics if light evoked response, the low density and sparse distribution of the Campana cells in the retina will limit their temporal and spatial resolution for image forming vision. Therefore, it is plausible to assume that Campana cells might play a role in a non-image formation function in the retina.

Another organizing hallmark of parallel pathways of the vertebrate retina is the segregation of light increment and decrement into ON and OFF synaptic pathways, which is initiated at BCs (Hartline, 1938; Wässle et al., 2009; Masland, 2012). In the inner retina, ON and OFF BCs transmit the increment and decrement luminance signals into ON and OFF RGCs by ramifying their axonal terminals into the ON and OFF sublaminae of the IPL, where they selectively synapse with ON and OFF neurons, respectively (Wässle et al., 2009; Wei and Feller, 2011; Masland, 2012; Sanes and Masland, 2015). However, Campana cells ramify their axonal terminals throughout the entire thickness of the IPL, and form synapses with both ON and OFF neurons, and likely transmit the ON signal, thus crossing the ON and OFF pathways. J-RGCs, which appear to have connections to Campana cells (data not shown), are considered OFF-RGCs with dendrites restricted to the OFF sublamina of the IPL. However, J-RGCs have been shown to have an ON-light response with wide field light stimulations (Kim et al., 2008; Joesch and Meister, 2016); therefore, Campana cells may be providing this size-based ON-response. Furthermore, it appears that Campana cells could provide inhibitory feedback to OPL neurons through glycine release from its dendrites (Heinze et al., 2007). Campana cells are uniquely poised to provide an ON-response to OFF neurons in the IPL, as well as to provide inhibitory feedback within the OPL, thereby providing a local enhancement to their excitatory signal. This may enable them to provide priming (excitatory), or stabilizing (inhibitory) signal to neurons in the IPL following the onset of light. However, while we have preliminary results that show Campana cells have the protein machinery to release both neurotransmitters, this is indirect evidence. Direct stimulation of Campana cells, as well as recordings from their direct synaptic partners will be required to confirm this proposed mechanism.

Interestingly, Campana cells have a relatively low density of ribbon synapse at their axonal plexus in comparison with conventional BCs (Tsukamoto and Omi, 2017) although they have a relatively high density of axonal terminals. Because these axonal terminals are filled with GlyT1, VGAT and SV2 labeled synaptic vesicles, it enables these Campana cells to release glycine into both ON and OFF pathways. This is similar to the dual transmitter releasing ACs, the starburst ACs, which release both Ach and GABA into both ON and OFF pathways. This is done through an intracellular system that involves both synaptic and calcium dependent regulation for modulating transmitter release (Yoshida et al., 2001; Lee et al., 2010). In future experiments, we will seek to find out if Campana cells use a similar mechanism. Additionally, it has been reported that light evoked ON responses can be transmitted to OFF pathways as both excitatory and inhibitory synaptic inputs through a “cross talk” circuit (Pang et al., 2007; Zhang and McCall, 2012). This is believed to only occur through amacrine cells, primarily Aii-ACs, and plays a significant role in the rod BC integration into the retinal circuit (Strettoi et al., 1994; Graydon et al., 2018). We are keen to determine whether Campana cells play any role in this “cross talk” circuit. We will seek to explore these possibilities in future studies.

## Acknowledgments

NIH grants R01EY012345 (N.T.), T32EY024234 (B.K.Y.), EY014800 (The Department of Ophthalmology and Visual Sciences of the University of Utah), Shared instrument grants: S10OD018033, S10OD023469 (Single Cell Genomics Core at BCM), P30EY002520 (R.C.). HHMI (K.D.), Research to Prevent Blindness (RPB, NY, New York): Department of Ophthalmology of Wayne State University School of Medicine (T.G.), and The Department of Ophthalmology and Visual Sciences of the University of Utah. The Ligon Research Center of Vision, Kresge Eye Institute, Dryer Foundation (T.G.). We want to thank Dr. Joshua Sanes for the BD-RGC mouse line, Dr. Alessandra Angelucci, for the primate retinal tissue, Dr. Botir T. Sagdullaev for the anti-GlyT1, Dr. Catherine W. Morgans for the anti-mGluR6, and Dr. Anand Swaroop for the anti-REEP6. We want to thank Dr. Zhuo-Hua Pan for the AAV2-GFP virus and the AAV2-GCaMP6m viral construct. Additionally, we want to thank Scott Gregory for assistance with video editing and production.

---

**Figure 1—supplemental video 1: Video of the 3D reconstruction of a Type 5 cone BC, Campana cell, and an Aii-AC.**

**This is associated with Figure 1C-1E**.

The three-dimensional rotating view of a type 5 cone BC (left; blue: cone photoreceptors), Campana cell presented in Figure 2D (middle; blue: DAPI), and an Aii-AC (right; blue: cone photoreceptors) labeled using AAV2-GFP to show the similarities and differences of the cell types. Background cells are colored in orange for clarity.

**Figure 2—supplemental video 1: Image stack of AAV2-GFP labeled Campana cell with anti-Cone Arrestin (C. Arr.), CtBP2, and mGluR6 staining.**

**This is associated with** **Figure 2A**.

Image stack of the same area shown in Figure 2A. A “*” indicates the soma of the Campana cell.

**Figure 2—supplemental video 2: Video of the 3D reconstruction of a Campana cell to photoreceptor synaptic connection.**

**This is associated with Figure 2B.**

A rotating view of the 3D reconstruction of the Campana cell to cone synaptic connection from a super-resolution image stack shown in Figure 3B. The Campana cell dendrite is green, the mGluR6 receptor is red, ribbon synapse is white, and the cone photoreceptor is blue.

**Figure 2—supplemental video 3: Video of the 3D reconstruction of Campana cell axonal ribbon synapse.**

**This is associated with Figure 2C and 2F.**

A rotating view of the three-dimensional reconstruction of the Campana cell axonal ribbon synapses from an image stack presented in Figure 3C to illustrate the co-localization of ribbons (white) inside the axonal varicosities of the Campana cell axonal terminals (green). This movie also shows a demonstration for how we determine ribbon synapses that are encased entirely within the cell in three dimensions.

**Figure 3—supplement figure 1.**
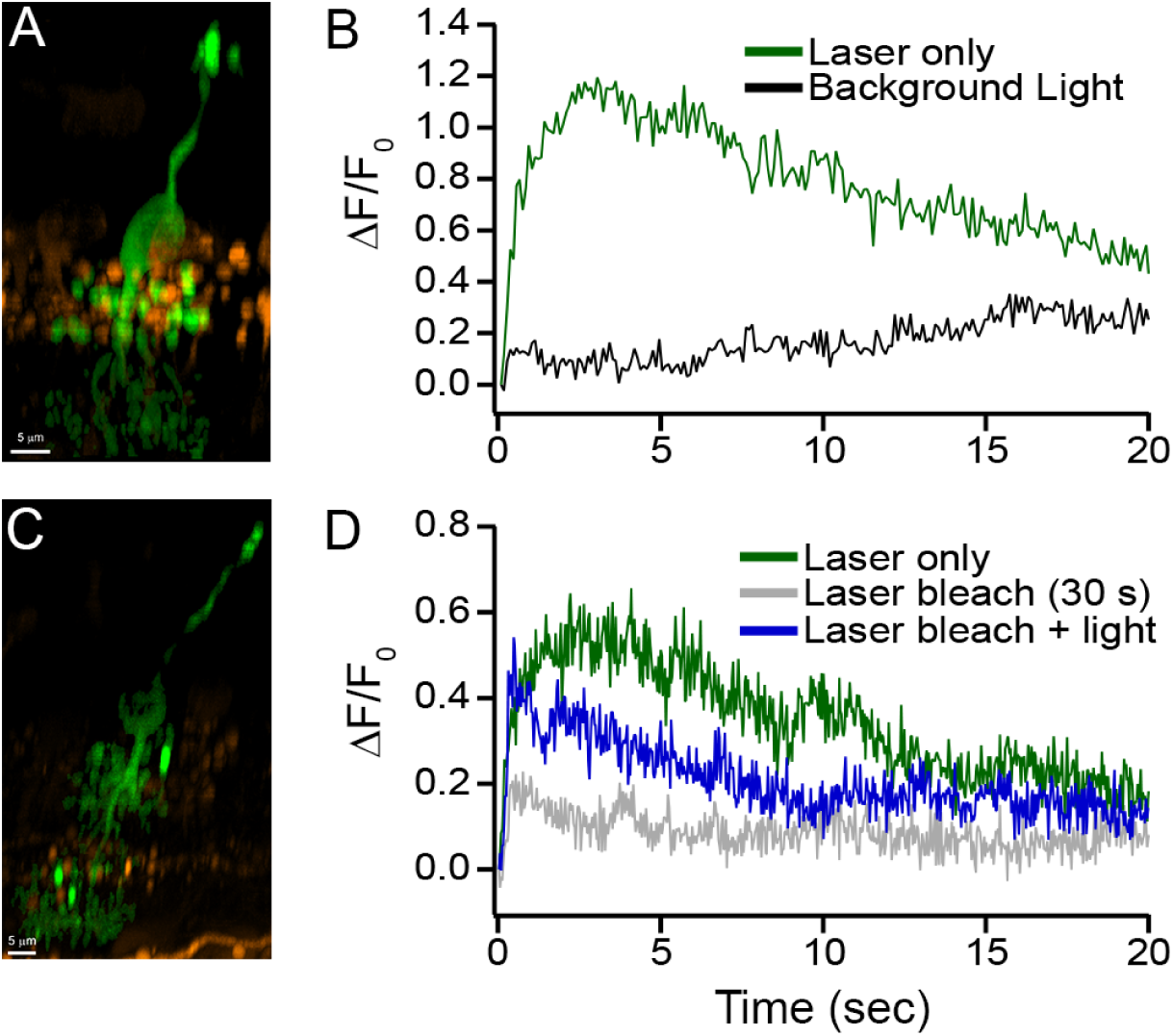
Campana cells show an increase in calcium concentration in response to two-photon laser scanning that is likely due to rod input. **This is associated with Figure 3.** **A:** A side view of the GCaMP6m fluorescence image of a Campana cell that had light responses recorded from its soma as shown in panel B. The cell was masked and isolated (green) from other cells (orange) for clarity. **B:** Change in GCaMP6m fluorescence intensity as a function of time during the two-photon scanning (green, laser only), and scanning by the two-photon laser after 10 minutes of green background light (black, background). **C:** A side view of another Campana cell that had light responses recorded from its soma, as shown in panel D. The cell was masked and isolated (green) from other cells (orange) for clarity. **D:** Change in GCaMP6m fluorescence intensity as a function of time in response to the two-photon laser scanning (green, laser only), scanning after a continuous 30 seconds laser scanning (grey, laser bleach), and a 10 ms UV LED light flash-evoked response after 30 seconds of continuous laser scanning (blue, laser bleach + light).

**Figure 5—supplement figure 1.**
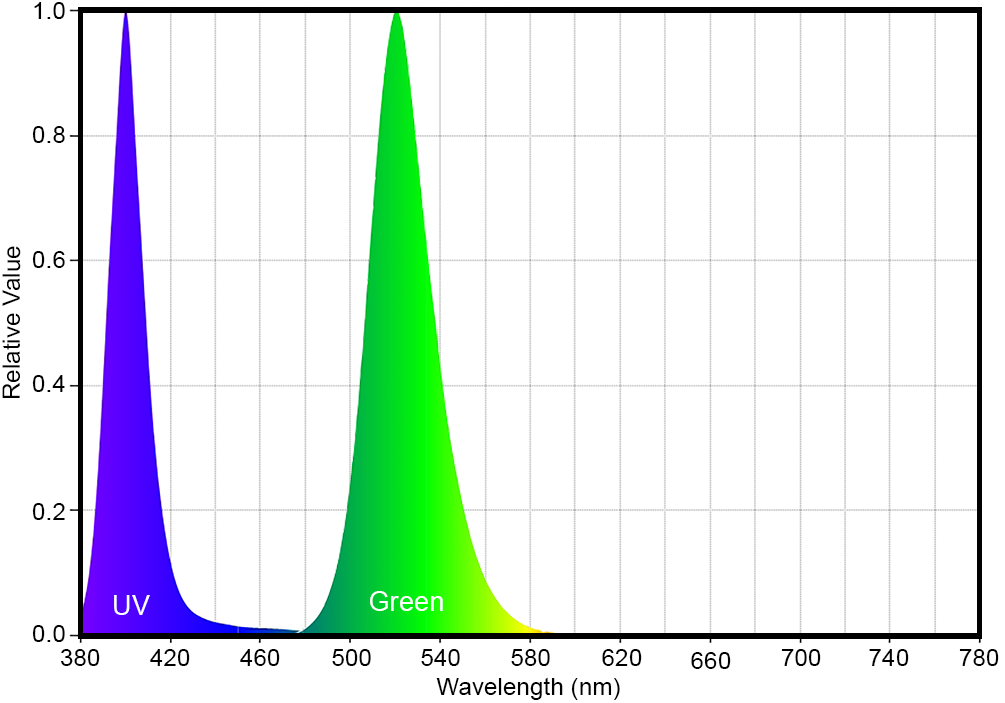
UV and green light color spectrum. **This is associated with Figure 5.** Spectrum measurements for the UV and green LEDs used in the experiments.

**Figure 6—supplemental video 1: Image stack of AAV2-GFP labeled Aii-AC and Campana cell with anti-GlyT1 staining.**

**This is associated with Figure 6C.**

Image stack of the same area shown in Figure 6C1. A “*” indicates the soma of the Campana cell.

